# Episodic memory retrieval success is associated with rapid replay of episode content

**DOI:** 10.1101/758185

**Authors:** G. Elliott Wimmer, Yunzhe Liu, Neža Vehar, Timothy E.J. Behrens, Raymond J. Dolan

## Abstract

Memory for everyday experience shapes our representation of the structure of the world, while retrieval of these experiences is fundamental for informing our future decisions. The fine-grained neurophysiological mechanisms that support such retrieval are largely unknown. We studied participants who first experienced, without repetition, unique multi-component episodes. One day later, they engaged in cued retrieval of these episodes whilst undergoing magnetoencephalography (MEG). By decoding individual episode elements, we found that trial-by-trial successful retrieval was supported by sequential replay of episode elements, with a temporal compression factor greater than 60. The direction of replay supporting this retrieval, either backward or forward, depended on whether a participant’s goal was to retrieve elements of an episode that followed or preceded a retrieval cue, respectively. This sequential replay was weaker in very high performing participants, where instead we found evidence for simultaneous clustered reactivation. Our results demonstrate that memory-mediated decisions are supported by a rapid replay mechanism that can flexibly shift in direction in response to task requirements.

**One Sentence Summary:** Recall of extended episodes of experience is supported by compressed replay of memory elements that flexibly changes direction depending on task temporal orientation.

## Main Text

Although a subject of intense study, the fine-grained mechanisms underlying how we retrieve episodes of experience are unknown (1). Understanding the supporting neurophysiological processes can reveal how episodes are represented in memory, and how they are subsequently retrieved to guide behavior (2, 3). Here we investigate whether episodes of experience are represented in a way that yields compressed sequential replay at retrieval, whether such replay supports successful retrieval, and whether the directionality of replay is flexibly tuned by internal goals.

Observations from animal studies have identified offline reactivation of sequences of hippocampal place cells that reflect past and future trajectories, thought to support memory consolidation, retrieval, and planning (4–6). Recently, animal studies have established a relationship between such replay strength and successful performance on spatial navigation tasks (4, 5). It has also been speculated that compressed replay might also support episodic memory retrieval in humans (7).

Human neuroimaging studies provide evidence for rapid cue-elicited reactivation of stimulus associations at retrieval (8–17) including overlapping replay of episode elements (18). A limitation of these studies is their inability to probe mechanisms supporting structured and temporally compressed reactivation, i.e. replay that proceeds at a rate faster than the original experience. An advance in human neuroimaging research has been a recent identification of rapid sequential replay of internal state representations (19, 20). Here, we leverage these same methods to ask whether sequential replay supports memory based decisions in humans.

We tested a hypothesis that episodic memory retrieval depends on rapid compressed replay of memory elements. Previous research demonstrating replay, which did not link replay to behavior, identified a short 40-50 ms lag between states (elements of a sequence) either during tasks involving lengthy planning periods or during undemanding rest periods (19, 20). Under similar conditions in rodents replay is known to occur preferentially during brief high-frequency sharp-wave ripple (SWR) events in the hippocampus (21–23). In contrast, theta-related sequence events are observed during active navigation and decision making (21, 22, 24, 25). The latter led us to expect that, during active memory retrieval, performance would be supported by replay events with a different and potentially longer lag between states.

Replay direction, forward or backward, is not always associated with particular task requirements in rodent research, though some studies show it is influenced by conditions such as active movement and reward receipt (20, 26, 27), potentially serving different computational functions (28). Recent MEG studies in humans have found reverse direction replay (19), or both forward and reverse replay (20). Based on these observations we expected replay direction would change flexibly based on internal states or task demands. In relation to our study design, we predicted replay would switch direction depending on whether the current goal was to retrieve memory components that followed a cued element, compared to having to retrieve memory components that preceded a cued element. In humans, replay onset has been associated with high-frequency power increases in the medial temporal lobe (MTL) (MTL) (20), and while we did not expect similar high-frequency changes, we nevertheless expected that the onset of memory replay events, irrespective of directionality, would be coupled to increased power in the medial temporal lobe (MTL).

Further, we reasoned that the strength of encoding, as reflected in better memory performance, would relate to enhanced memory consolidation (1, 7). Greater experience is associated with less marked replay in rodents (25, 29), and this predicts a less dominant expression of replay in participants who show near-ceiling memory performance. In these participants, theoretical proposals suggest a form of clustered pattern completion for episode elements (9, 30, 31). Importantly, this predicts that, within participants, trial-by-trial sequenceness strength should relate positively to trial-by-trial retrieval success. At the same time, if very high performing participants do not rely to the same degree on a replay mechanism for retrieval, then across participants this entails that mean sequenceness strength could be negatively related to mean memory performance.

We designed a novel episodic memory task and combined this with our recently-developed MEG analytic methods (19, 20). In brief, on day 1 participants experienced temporally extended self-oriented episodes, where each single-exposure episode was itself composed of five discrete and unique picture stimuli that were assembled into a narrative story (**Fig. 1a** and **Fig S1**). Following an overnight consolidation period, we then elicited cued retrieval of these episodes whilst obtaining MEG data to index fast neural dynamics supporting retrieval (**Fig. 1b**).

**Fig. 1.**
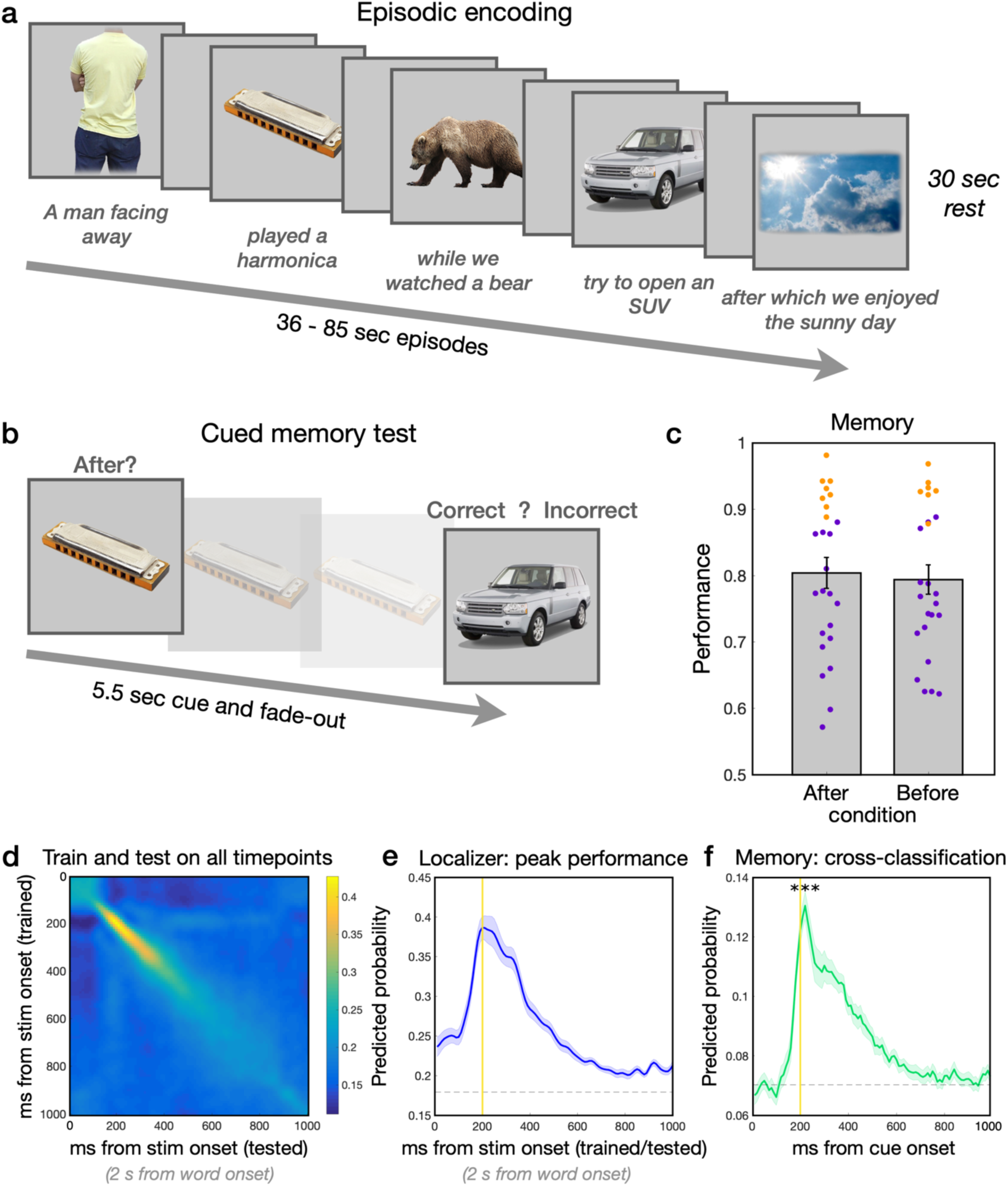
Experimental design and decoding of the episode elements. (**a**) On day 1, in the episodic encoding phase we presented subjects with eight extended non-spatial episodes, with a single exposure per episode. Episodes contained five stimulus elements. The first four episode elements were selected from six distinct picture categories. Participants were incentivised to encode the precise order of the episode elements. (**b**) On day 2, in the episodic memory test phase, participants retrieved episodes in two conditions. In the ‘after’ condition, participants were asked to identify whether a subsequent probe element came after the cue element. Following a 5.5 s retrieval period, a test probe presented. The sequential order referred to any stimulus from the same episode that followed this cue; here, the depicted answer would be ‘correct’. By contrast, in the ‘before’ condition, participants were asked to identify whether a subsequent probe element came before the cue element. (**c**) Mean memory performance in the after and before conditions. Purple dots represent individual data points for regular performance participants with sufficient incorrect response (error) trials free from MEG artifacts for accuracy analyses (after, *n = 17*; before, *n = 18*); the remaining very high performance participants are shown in orange (see also **Fig. S1**). (**d**) Classifier performance for episode element categories presented during the localizer phase, training and testing at all time points, showing good discrimination of the 6 categories used to compose the first four episode elements. In localizer trials, note that a word naming the upcoming stimulus appeared 2 s before the stimulus, contributing to above-chance classification at 0 ms. (**e**) Peak classifier performance at 200 ms after stimulus onset in the localizer phase (depicting the diagonal extracted from panel d; see also **Fig. S3**). Dashed line represents the mean across time 95 % level of randomly shuffled classifier labels. (**f**) Application of the trained classifier (at 200 ms) to cue onset in memory retrieval trials demonstrated above chance decoding of the current on-screen category during retrieval. Dashed line represents the maximum value of classifier during pre-trial baseline; performance was compared to this baseline value. (Error bars and shaded error margins represent standard error of the mean (SEM).)

As a first step we confirmed we could reliably identify neural patterns associated with individual episode elements, each drawn from one of six different stimulus categories. Note that the final element of each episode was not taken from a decoded category. A classifier trained on the localizer phase showed successful discrimination of the categories that made up the episodes with peak decoding at 200 ms after stimulus onset (**Fig. 1d-e**; **Fig. S3-S4**), in line with previous reports (19, 20). In an exploratory low-powered analysis of single stimuli, we found that these categories were also evident as clusters of similarity in trained sensory weights (**Fig. S3**; Supp. Results). The trained classifier generalized to the memory retrieval phase, showing significant across-phase classification of cue category (peaking at 210 ms after the cue; compared to chance at 200 ± 10 ms (the peak timepoint in localizer phase) t_(24)_ = 9.80, p < 0.001; **Fig. 1f**).

To test specific predictions of a replay mechanism underlying episodic retrieval, we next sought evidence for compressed sequential reactivation of episode elements during the retrieval period. In this analysis, we first derived measures of category evidence – representing reactivation of memory elements – at each timepoint by applying the trained classifiers to retrieval period MEG data. We then tested for lagged cross-correlations between episode element reactivations across the retrieval period, yielding a measure of ‘sequenceness’ in both forward and backward directions (19, 20) (**Fig. S2; Methods**). Following an approach used in previous reports, to identify time lags showing potential sequenceness and examine a relationship to individual differences in memory performance, we tested for a difference between forward and reverse direction components (19, 20). Our initial analyses focused on memory retrieval in the *after* condition. Here participants are asked to identify whether a probe element came sequentially after the cue element, a condition we considered would be easier and more naturalistic than the *before* condition.

For the individual differences analysis, we identified a state-to-state time lag of interest by focusing on correct trials, where we expected stronger sequenceness. In the *after* condition, we identified an overall dominance of reverse replay (backwards > forwards sequenceness) during correct trials, peaking between 100-120 ms (**Fig. 2a**). The peak lag between 100-120 used for the independent individual difference analyses does not survive correction for the number of tests across lags, so it should not be interpreted on their own. Of interest, this time window for rapid online retrieval represents a longer state-to-state time lag than the 40-50 ms lag found in other experiments reporting replay during extended planning or rest (19, 20). As in rodents, these fast resting replay events (with 40-50 ms state-to-state time lag) have been associated with sharp-wave ripples in humans (20). However, rodents also show sequence events during ongoing behaviour that are associated with ongoing hippocampal theta rhythms (24, 25), though heretofore such online sequence events have not yet been identified in humans.

**Fig. 2.**
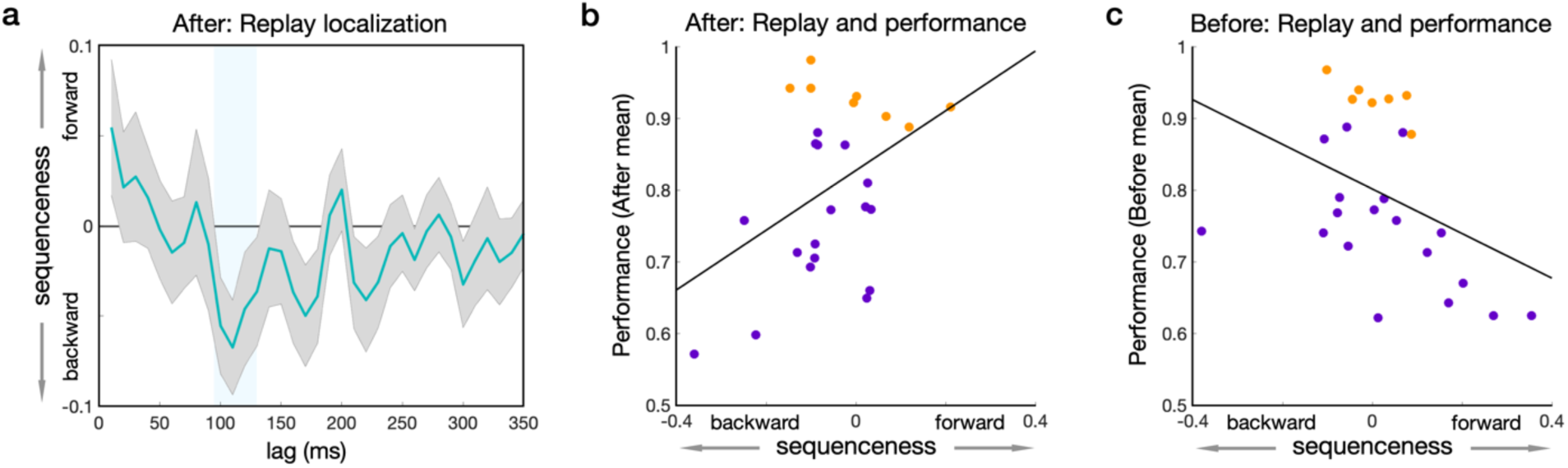
Mean sequenceness (replay) in the *after* condition and the relationship between sequenceness and performance in the *after* and *before* memory retrieval conditions respectively. (**a**) In the *after* condition, mean forward minus backwards sequenceness for correct memory trials (when participants accurately answered the memory question). On correct trials, a peak of reverse sequenceness was observed at lags from 100-120 ms. This time window was used for subsequent analyses. (Shaded error margins represent SEM.) (**b**) In the *after* condition, stronger mean reverse sequenceness on correct trials correlated negatively with overall mean memory performance (percentage of correct trials). As in Fig. 1c the data points for the regular performance participants are shown in purple; high performance participants are shown in orange. (**c**) In the *before* condition, stronger forward sequenceness related to lower performance. The overall results in the *after* and *before* conditions support a stronger role for replay in retrieving weaker memory traces.

To provide an initial test of an association between replay and episodic retrieval, we examined the relationship between replay strength in correct trials and overall memory performance. We found that differential sequenceness correlated with mean memory performance (100-120 ms lag; r = 0.4254, p = 0.034; **Fig. 2b**). As sequenceness was on average negative – showing predominantly a reverse direction of replay – this suggests that stronger reverse replay is a characteristic of individuals with weaker performance. Notably, this relationship between replay and memory strength is in line with the findings in rodents showing stronger sequenceness during initial acquisition compared to later high performance (25, 29).

As an initial test of our prediction that internal goals – whether looking forward or backward in time through an experience – are important for retrieval and replay, we examined whether the relationship between replay and individual differences in performance changed from the *after* compared to the *before* condition. If task goal affected replay, we would expect stronger forward sequenceness to be related to weaker performance. Indeed, in the before condition we found the degree of dominantly forward sequenceness correlated negatively with mean memory performance (100-120 ms lag; r = −0.4077; p = 0.0431; **Fig. 2c**). We then examined the strength of these correlations using a conservative permutation approach, where the goal was to test for a potential influence of a decreasing number of correct trials per mean datapoint going from high to low performing participants. In the after condition the correlation between sequenceness and mean performance exceeded the conservative permutation threshold (adjusted 5 % level 0.041, versus p = 0.034) while the strength of the before condition effect fell just outside the permutation threshold (adjusted 5 % level 0.0395, versus p = 0.043).

Comparing the after and before results, we found that the correlation between sequenceness and performance in the after condition differed significantly from that in the before condition (z = 2.411; p = 0.0159; two-tailed, conservatively using the test for dependent correlations). This provides initial support for our prediction that retrieval orientation influences the characteristics of replay that support behaviour. Importantly, the results in the after and before conditions both indicated that replay was stronger in participants with lower overall performance, with replay playing a lesser role in retrieval for participants with near-ceiling levels of performance. However, these results do not indicate per se whether sequenceness is positively or negatively related to trial-by-trial retrieval success.

We found no across-participant relationship between mean sequenceness and behavior in the shorter 40-50 ms state-to-state time lag as identified in previous studies (**Fig. S4**). In an exploratory analysis that examined evidence for sequences of episode elements present in any of the other 7 episodes (but not present in the current episode), we found a numerically negative sequenceness effect at 40 ms, but again found no relationship to memory performance (**Fig. S4**).

We next exploited analytic techniques that simultaneously examined the influence of forward and backward sequenceness on memory performance. First, we examined the relationship between sequenceness and individual differences in performance. This confirmed the above results, namely that weaker memory performance related to stronger reverse replay in the after condition, and to stronger forward replay in the before condition (see **Supp. Results**).

To examine whether trial-by-trial forward and reverse sequenceness related positively or negatively to retrieval success, we utilized multilevel regression analyses. These analyses include more than a hundred datapoints per participant and are thus the most highly powered analyses in the current experiment. For these analyses we excluded very high performing participants, as they have too few incorrect trials to support reliable estimates. We first independently localized a time lag of interest using leave-one-participant-out cross-validation procedure, again identifying a peak time lag of 110 ms in all participants except one very high performing participant (who showed a lag of 170 ms); thus, we used a 110 ± 10 ms lag for all regular performance participants with sufficient incorrect trials for analysis.

In the after condition, we found that reverse sequenceness from 100-120 ms related positively to trial-by-trial retrieval success (multilevel regression on accuracy in n = 17 participants with sufficient incorrect trials; forward *β* = −0.1336 [−0.299 −0.020]; z = −1.714; p = 0.0920; reverse *β* 0.1881 [0.042 0.338]; z = 2.416; p = 0.0176; **Fig. 3a**). An example of a reverse sequence in the *after* condition for a single participant is shown in **Fig. 3c**. By contrast, in the before condition forward, but not reverse, sequenceness related positively to accuracy (regression in n = 18 participants with sufficient incorrect trials; forward *β* = 0.160 [0.014 0.305]; z = 2.202; p = 0.0264; reverse *β* = −0.0564 [−0.207 0.091]; z = −0.763; p = 0.470; **Fig. 3A**). An example of a forward sequence in the before condition for a single participant is shown in **Fig. 3d**. In the after and before conditions, the forward or reverse direction of sequenceness that related to trial-by-trial retrieval success was the same as the performance-related direction identified in the individual difference analyses. The same relationships between sequenceness and retrieval success were also found in models where we included all participants (**Table S3**). Importantly, we found a significant interaction between both forward and reverse replay and the after versus before goal condition (condition by forward replay *β* = −0.1608 [−0.271 −0.049]; z = −2.865; p = 0.0032; condition by reverse replay *β* = 0.1408 [0.029 0.250]; z = 2.499; p = 0.0096; **Fig. 3b**; n = 15 participants with sufficient incorrect trials in both the after and before conditions).

**Fig. 3.**
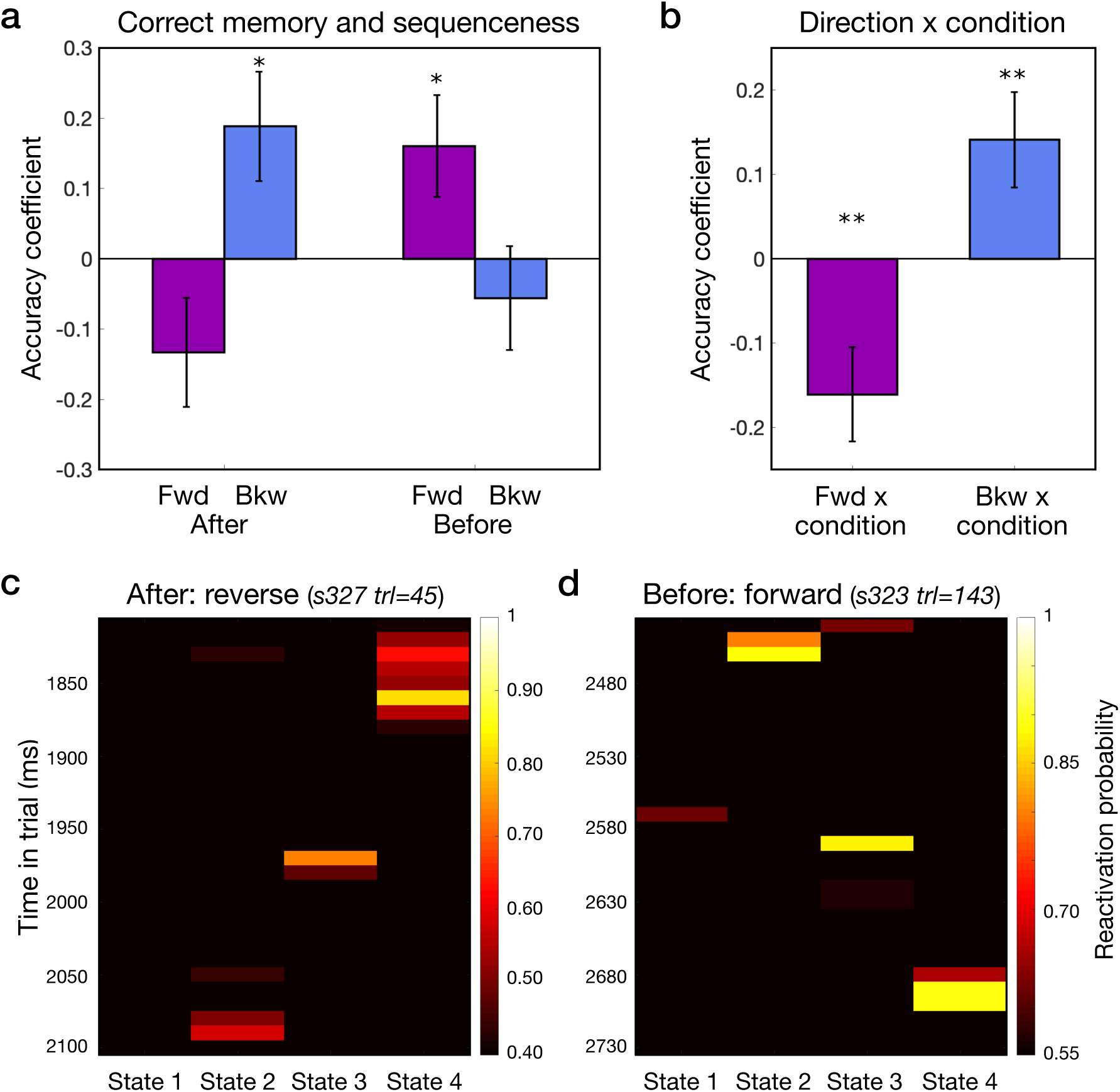
Relationship between forward and backward sequenceness and trial-by-trial memory retrieval success in the after and before conditions. (**a**) In the *after* condition (left), successful memory retrieval was supported by reverse sequenceness. In the *before* condition (right), retrieval was supported by forward sequenceness. See also **Fig. S6 for** individual participant regression coefficients derived from a single level analysis. (**b**) Interaction of replay direction (forward, backward) by condition (after, before) showing a stronger effect of forward replay on trial-by-trial successful memory retrieval in the before condition, and a stronger effect of backward replay on successful memory retrieval in the after condition. (The regular performance group in the combined sequenceness analysis included n = 15 participants common to the regular performance group across the after and before conditions.) (**c**) Example of reverse sequenceness in the after condition. (**d**) Example of forward sequenceness in the before condition. (s = participant; trl = trial; *p < 0.05; **p < 0.01; error bars represent SEM).

As in the individual differences analyses, in the trial-by-trial analyses, we did not find any relationship between the sequenceness measure derived from the alternative 7 episodes (‘other episode’ sequenceness) and retrieval success at a 40-50 ms lag (identified via leave-one-out cross-validation on this sequenceness measure; p-values > 0.35; **Supp. Results**; **Fig. S6**; **Table S5**), while sequenceness derived from the current episode transitions remained significant. An additional other episode sequenceness measure derived from a 100-120 ms lag was also not related to behavior (**Fig. S6**;**Table S6**).

It is possible that the underlying representations of episodes may change across the many cued retrieval events, despite the original episodes not being actually re-experienced. To investigate this possibility, and in particular whether our results were driven by effects that appear after extensive practice, we examined whether the sequenceness-accuracy relationship changed over the course of the retrieval task. We found, if anything, a tendency for a numerical decrease in the sequenceness-accuracy relationship over the course of the experiment, and this was true for both the after and before conditions (**Table S3**). It is also possible that participants developed strategies to sequentially reactivate items in different orders with respect to the after and before conditions. However, we found no evidence for this in participant self-reports (**Table S2**).

In a final control analysis, we address concerns about potential bias in our analyses. Thus, we conducted simulations of the full processing and analysis pathway, from the generation of localizer data through to the final step of multilevel regressions that relate sequenceness to retrieval success. The simulation results confirmed that the relationship between randomly generated MEG data and behavioral measures was what would expected by chance: the false positive rate was near an expected 5 % level in both the after condition (0.055) and before condition (0.04; **Fig. S7**).

The relationship between sequenceness and successful memory retrieval in both the after and before conditions provides a clear link between sequenceness and behavior. While the initial individual differences analyses found relatively stronger sequenceness in regular performing participants, these trial-by-trial results demonstrate that within regular performance participants, sequenceness strength is positively related to retrieval success. Incorporating the results of the individual difference analyses and the trial-by-trial analyses, we establish a double dissociation between replay direction and a participant’s internal goal condition during retrieval. These findings demonstrate a flexibility in replay directionality that goes beyond previously reported effects of external events such as reward receipt (20, 27).

Inspired by neurophysiological studies showing that the hippocampus is a source for replay events, we next examined whether replay event onset related to power increases within the medial temporal lobe (20). Candidate replay onsets were identified by locating sequential reactivation events showing a 110 ms lag, applying a stringent threshold to these events, and using beamforming analysis to localize broadband 1-45 Hz power changes related to replay event onsets. For reverse replay events (in the *after* condition) and for forward replay events (in the *before* condition), this analysis localized activity at replay onset to a region of right anterior MTL, encompassing the hippocampus and entorhinal cortex (after: z = 3.72, p <0.001 whole-brain FWE; before: z = 3.73, p < 0.001 whole-brain FWE; **Fig. 4a-b**; **Table S2**), consistent with human fMRI results during rest in a cognitive paradigm (32). The increase in MTL power was selective to replay onset, with an additional secondary peak in the after condition 1 lag later at 110 ms (**Fig. S8**). In the after condition, replay onset also related to activity in two significant clusters in the right visual cortex (**Fig. S8**; **Table S2**). Finally, we found evidence for stronger power immediately preceding replay onset in the left anterior MTL in participants with lower performance (z = 3.82, p = 0.003 whole-brain FWE; **Fig. S8**).

**Fig. 4.**
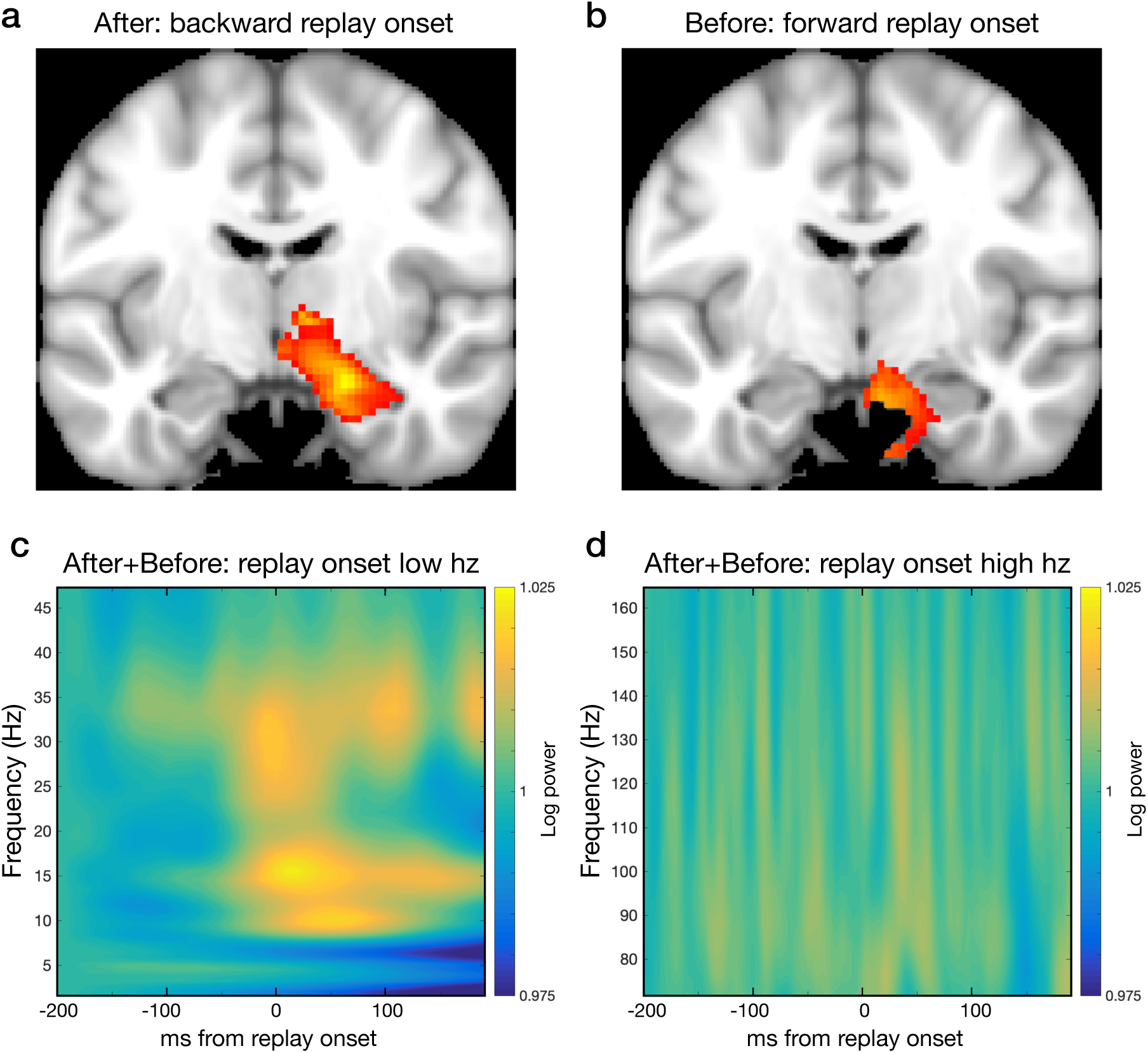
Beamforming analysis of power increases at the onset of sequenceness events and time-frequency analyses of replay onset. (**a**) In the after condition, power in the right anterior MTL increased at onset of reverse sequenceness events (n = 25). (**b**) In the before condition, power in the right anterior MTL increased at the onset of forward sequenceness events (n = 25). (Statistical maps thresholded at p < 0.001 uncorrected, for display; for unthresholded statistical maps see: https://neurovault.org/collections/6088/) (**c**) Time-frequency analysis showing power change relative to replay onset across the after and before conditions in frequencies up to 50 Hz. 0 ms represents the onset of putative replay events. (**d**) Time-frequency analysis of replay onset showing no high frequency power change relative to replay onset across the after and before conditions (using data sampled at 600 Hz).

Replay onset was associated with broadband power increases from approximately 8 Hz up to 45 Hz in across the after and before conditions (**Fig. 4c** and **Fig. S9**). In the frequency range of our element-to-element lag (8-12 Hz, approximately human alpha), we found an increase in power at replay onset (t(24) = 4.267 [0.003 0.008], p < 0.001). However, we found no evidence for power increases in the high gamma frequency range that have been associated with replay events during rest (events that may be related to sharp-wave-ripple events; 120-150 Hz; t(24) = 1.150 [-0.001 0.005], p = 0.262) (20) (**Fig. 4d** and **Fig. S9**).

Finally, as very high performing participants did not show any relationship between replay and performance, we examined the hypothesis that retrieval for strongly encoded memories is based on clustered pattern completion. Across all participants, with a rapid appearance following cue onset, we found significant evidence for reactivation of within-episode elements compared to other-episode elements, none of which were displayed on the screen (average across timepoints showing the strongest classification of on-screen cues, 210 ±10 ms post-cue t_(24)_ = 3.978, p < 0.001; **Fig. 5a**). A reactivation event from a single participant is shown in **Fig. 5b**.

**Fig. 5.**
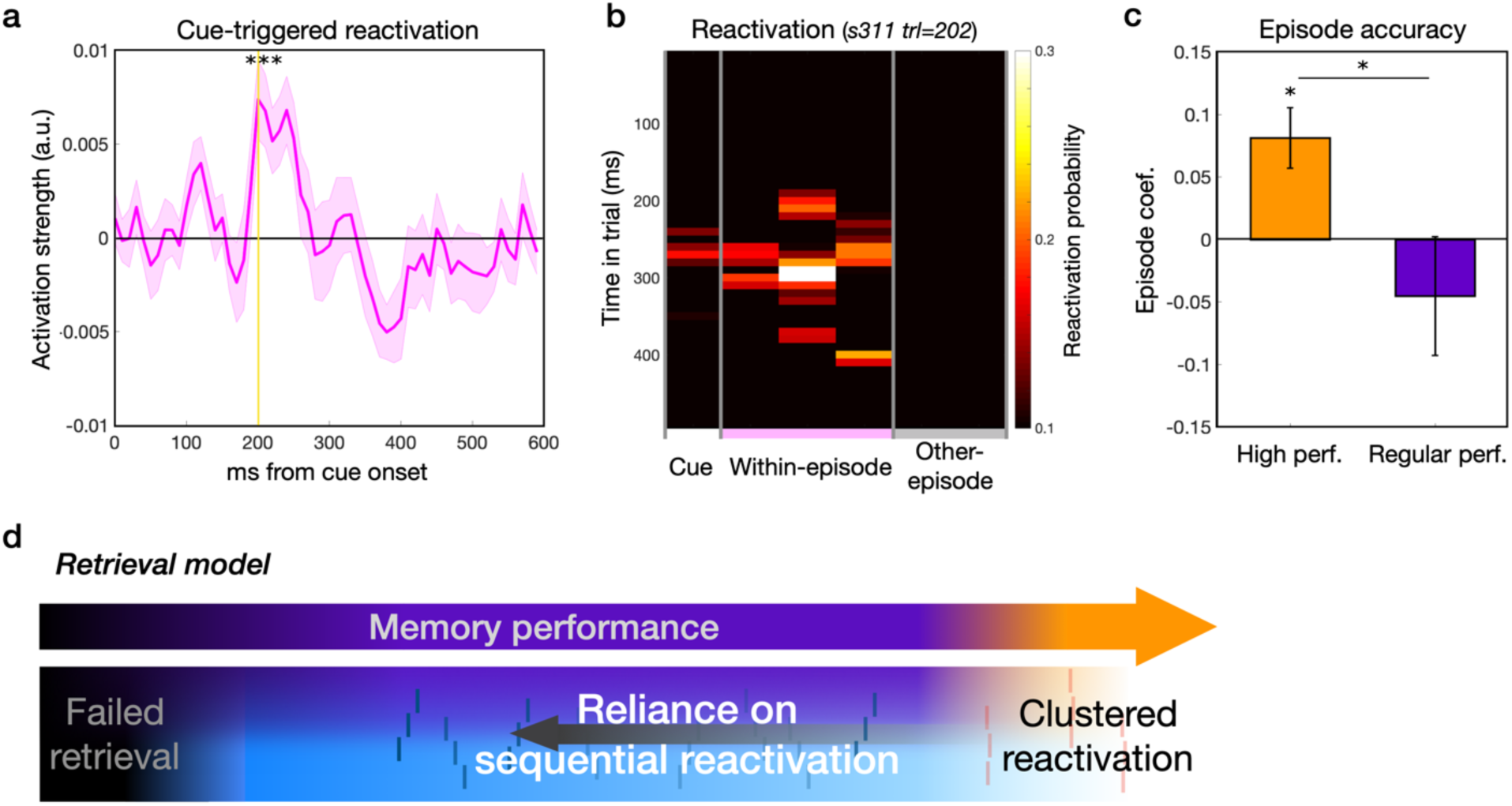
The relationship between cue-evoked reactivation and performance. (**a**) Across both the *afte*r and *before* conditions, we found evidence for cue-evoked reactivation of the elements present in the episode, peaking 200-250 ms after cue onset. (Shaded error margins represent SEM.) (**b**) Example of cue-evoked reactivation of within-episode elements in a single trial in a single participant. (**c**) Cue-evoked reactivation related to mean performance in a given episode for the high performance participants, but not regular performance participants (group breakdown based on number of incorrect trials across both the after and before conditions; high *n = 10*; regular *n = 15*; error bars represent standard error.) (**d**) Retrieval model illustrating the relationship between memory and sequenceness across- and within-participants. Across participants, higher mean memory performance was associated with weaker sequenceness and stronger cue-evoked reactivation of episode elements (‘clustered retrieval’). Within-participants, in regular performing participants, stronger trial-by-trial sequenceness positively related to trial-by-trial retrieval success.

To examine the relationship between the cue-evoked reactivation effect and memory in very high performance participants, instead of a contrast of correct versus incorrect trials, we used a measure of mean performance for the episode cued on the current trial (a graded measure from 0 to 1). Cue-evoked reactivation was averaged across the 200-250 ms peak difference of current versus other episode elements. Reactivation positively related to performance on a given episode in very high performing participants (n = 10; *β* = 0.0795 [0.0321 0.1250]; t = 3.442; p < 0.0008; **Fig. 5c**), an effect stronger in high compared to regular performance participants (regular *β* = −0.0440 [−0.1387 0.0512]; t = −0.918; p = 0.3568; difference *β* = 0.125 [0.005 0.244]; t = 2.035; p = 0.0376; **Fig. 5c**). In a follow-up analysis, we found that the effect in very high performance participants related positively to evidence for within-episode elements (p < 0.04) and related negatively to evidence for other-episode elements (p < 0.06); neither measure related to accuracy in the regular performing participants (p-values > 0.29).

Additionally, although based on a very low number of trials, in very high performing participants we found that correct trials related to higher cue-evoked reactivation as compared to incorrect trials (*β* = 2.497 [0.4009 4.444]; t = 2.409; p = 0.024; **Fig. S10**). We found no significant relationship between cue-evoked responses and accuracy in regular performance participants (regular *β* = 0.6291 [-0.3483 1.6681]; t = 1.281; p = 0.2072; difference *β* = 1.872 [-0.052 4.385]; t = 1.633; p = 0.116). Importantly, in regular performing participants, the trial-by-trial relationship between sequenceness and accuracy in both the after and before conditions remained significant when including cue-evoked reactivation in the same model while the cue-evoked reactivation measure was not significant (**Fig. S10**; **Table S5**).

In additional control analyses, we examined the relationship between memory and responses to the cued category itself as well as overall classifier strength throughout the remainder of the retrieval period. First, responses to the cued element on the screen did not relate to mean episode performance or accuracy across conditions (from 200-250 ms; p-values > 0.13). However, in regular performing participants in the after condition (where all cues are decodable) a positive relationship was evident between accuracy and cue responses (p = 0.0016; **Fig. S10**; **Table S6**). Importantly, however, the relationship between backward sequenceness and memory remained significant in a model that included cued category responses, suggesting potential independent mechanisms contributing to performance. We found no relationship between responses to the cued category and forward or backward sequenceness itself in either the after or before conditions (p-values > 0.42).

In the post-cue retrieval period (following the initial 200-250 ms cue-evoked response period), we found no relationship between successful retrieval and classifier evidence for the on-screen cue stimulus, the within-episode categories, or the other episode categories (on average across the remaining 250 – 3670 ms time window, p-values > 0.19). The classifier results also did not show differential evidence for the fading cue: we found no overall difference in classifier evidence between the cued on-screen stimulus, the within-episode categories, and other episode categories (p-values > 0.83). Finally, in an exploratory analysis of simultaneous joint reactivation of different categories, while we could identify putative simultaneous reactivation events during the retrieval period, we found no relationship between these events and performance in regular performance participants (**Fig. S10**), supporting the importance of sequential reactivation for successful episodic memory retrieval.

During episodic memory retrieval in humans, we show that a rapid sequential replay of episode elements relates to differences in memory performance. Our primary finding is a demonstration that stronger trial-by-trial sequenceness relates to retrieval success across conditions. Across-participants, we found that regular performance participants exhibited stronger sequenceness than high performance participants. As illustrated in the memory retrieval schematic (**Fig. 5d**), these results are complementary and a seeming contradiction is a reflection of Simpson’s paradox (33). Given the dominance of replay in regular performing participants, replay may play a functional role in “piecing together” individual retrieved elements. Additionally, we find that replay proceeds in the opposite direction to what might be expected, i.e. replay flows from distal episode elements to the proximal cued element (34). In general, our results indicate an important role for replay in online memory retrieval, with an element-to-element lag of 100-120 ms, establishing a novel connection between replay and ongoing behaviour in humans that has only recently been demonstrated in animal research (4, 5, 29).

Replay events spanned a temporal horizon of seconds during retrieval, in contrast to a single instance of clustered pattern completion (9, 30). The latter pattern characterised very high performing participants alone, where cue-evoked reactivation closely resembled pattern completion. We cannot exclude a possibility that an absence of sequential replay in very high performing participants might reflect a difficulty in detection, perhaps due to a sparse distribution or rapid decay of replay event frequency. Similarly, our results could be biased towards detecting stronger sequenceness in regular performing participants, who exhibit a stronger engagement of retrieval processes, which in turn could provide greater evidence for classification of sequential activation. Alternatively, when episodes are strongly encoded during an experience itself, different representations might begin to form, where retrieved order information is no longer represented by sequential replay but instead by the clustered reactivation pattern we observed. A potentially related finding of a decreasing expression of replay with increasing experience has been reported in rodents (25, 29). Here we speculate that in high performing participants, episodes are more strongly encoded and potentially enhanced by spontaneous reactivation and replay during post-learning rest and sleep (6, 35–38), and these representations may be differentially supported by cortical systems (30, 31, 39, 40).

Replay in the current experiment showed an element-to-element lag of approximately 110 ms, representing a temporal compression factor of 60 to 150. This compression is in line with, or exceeds, the degree reported in offline place cell sequences in rodents (41, 42). Previous MEG research examining replay in humans report a shorter 40-50 ms lag between replayed elements for very well-learned sequences (19, 20). These studies allowed for tens of seconds of planning or involved acquisition over minutes of rest; further, replay during rest was related to putative SWR events (20). This contrasts with our current experiment where there was a requirement for relatively rapid ‘online’ decisions.

These different effects, influenced by task demands, parallel well-established results in animals. Thus, theta-related sequence events are found predominantly during active navigation, while replay events during high-frequency SWRs are found during rest and sleep (21–25). Based on a close association between animal and human replay during putative SWR events, as demonstrated recently (20), and the important distinction between the previous results pertaining to rest and current results that reflect active behavior, it is instructive to speculate on connections between our current findings and an expression of sequenceness observed in rodents, specifically that which relates to theta sequences. However, any suggested connection needs to be tempered by substantial differences between animal spatial navigation and human episodic memory. More extensive research is needed to fully explore any potential connection.

Episodic memory experimental designs utilizing actual extended sequences of experiences as episodes, instead of a more traditional use of multiple different static images, trade off benefits of increased ecological validity against a potential disadvantage of necessitating repeated testing of episodes. The use of repeated probes of episodes is often necessary when using decoding approaches, where the analyses require many exposures to the episode elements during training of a decoder. In some cases, experiments include re-exposures to the original episodes (18). As in real-life experiences outside of the lab, memory episodes in our experiment were experienced only a single time at encoding. Repeated testing on the other hand may alter the underlying memory trace or lead to increasing reliance on retrieval strategies, and we acknowledge this as an important caveat to studies of this type. Importantly, we found no change in the positive trial-by-trial sequenceness-memory relationship over the course of the experiment.

Individual episodes of experience are important building blocks for creating a representation of the structure of the world (2). Episodic representations that support replay are likely to be important for how we successfully navigate spatial, social, and abstract environments (3, 6, 43–47). In turn, memory closely interacts with decision making (e.g. 10, 46). The ability to reactivate episodes in a highly compressed manner provides a novel mechanism for very rapid retrieval and replay of previous experiences during choice (48–50), and our findings can motivate new directions of research into the relationship of memory encoding, consolidation and decision making. Further, the flexible direction of episodic retrieval replay events that we identify may affect choice dynamics. We speculate that sequential replay flexibility and strength might serve as markers for an impaired associative binding between memory elements caused by negative emotional events. Impaired, or pathologically disturbed, memory organization has a strong negative impact on well-being and behaviour, and future human research into memory replay might also provide novel insights into memory disturbances seen in psychiatric disorders such as post-traumatic stress disorder and schizophrenia (51, 52).

## Acknowledgments

The authors thank Zeb Kurth-Nelson for helpful discussions. This work was supported by a Wellcome Trust Investigator Award (098362/Z/12/Z) to R.J.D. Y.L. is supported by a UCL Graduate Research Scholarship and an Overseas Research Scholarship. The Max Planck University College London Centre is a joint initiative supported by University College London and the Max Planck Society. The Wellcome Centre for Human Neuroimaging is supported by core funding from the Wellcome Trust (203147/Z/16/Z).

## Author contributions

G.E.W., Y.L., and N.V. designed the experiment. G.E.W. and N.V. collected the data. G.E.W and Y.L. wrote the analysis code, analyzed, and interpreted the data. T.E.J.B. and R.D. contributed to data interpretation. G.E.W. wrote the paper with input from Y.L., N.V., T.E.J.B., and R.J.D.

## Competing interests

Authors declare no competing interests.

## Methods

Twenty-eight healthy volunteers participated and completed both sessions of the experiment. Participants were recruited from the UCL Institute of Cognitive Neuroscience Subject Database. Data from three participants were excluded due to poor memory performance (described below) leaving data from 25 participants for analyses (14 female; mean age 24 (range 18-32). Participants were required to meet the following criteria: age between 18-35, fluent English speaker, normal or corrected-to-normal vision, without current neurological or psychiatric disorders, no non-removable metal, and no participation in an MRI scan in the two days preceding the MEG session. The study was approved by the University College London Research Ethics Committee (Approval ID Number: 9929/002). All participants provided written informed consent before the experiment. Participants were paid for their time, for memory performance (up to £10 based on percent correct performance above chance), in addition to a bonus for localizer phase target detection performance (up to £2).

Participants were excluded from analysis if two of the following three criteria were met: (1) accuracy below 50 % on the cued retrieval task on the second day, (2) accuracy below 50 % in the episode component re-ordering task on the second day, and (3) indication on the post-experiment questionnaire that the participant had mentally reordered the episodes from their original day 1 order. As the MEG analyses tested for reactivation of sequences of episode elements based on the original order, relatively poor memory for the order of episode elements (in the post-test) and/or a report of re-ordering the episodes (in the post-questionnaire) were part of the exclusion criteria. As noted above, 3 participants from the initial 28 were excluded based on these criteria. In the current sample, no participants were excluded based on MEG decoding performance, specifically, the classification of the 6 categories in the MEG localizer phase data.

### Experimental Task

We designed our memory experiment to investigate the neural processes supporting retrieval of episodic experiences where the original episodes were only experienced once, similar to many experiences outside the lab; this is in explicit contrast to paradigms with many repetitions of the same (sequence of) stimuli. Retrieval was also separated from encoding by approximately 24 hours, again to increase ecological validity. Episodes were designed such that they were made up of elements from 6 different categories. In order to be able to classify many varying episode elements, without pushing the limits of pattern classification or participant alertness for long-duration scans, we designed our experiment using well-identified categories deployed in previous fMRI and MEG studies of memory and perception (e.g. 10, 12, 15, 52). Participants were explicitly instructed that memory episodes were made up of 6 categories of stimuli (faces, buildings, body parts, objects, animals, and cars), and then shown examples of these categories. Note that we did not expect participants to think of abstract category-level information during retrieval but instead expected participants to retrieve individual elements, without explicitly categorizing the items. We utilized categories of stimuli because we predicted that category-level information would provide the largest source of across-stimulus variability in neural responses.

On the first day of the experiment, in a testing room environment participants experienced 8 different temporally extended episodes with one single exposure per episode (**Fig. 1a**). Participants were told their performance on memory questions that tested their knowledge of the correct sequential order of stimuli would influence the amount of a monetary bonus. Episodes were composed of 5 discrete picture elements and an accompanying story written from a first-person perspective. On the second day, participants returned for an MEG scanning session where they completed a cued retrieval phase and a category localizer phase during the acquisition of MEG data (**Fig. 1b**). Behavioral piloting in a separate sample of participants was used to optimize the design and ensure that memory retrieval performance on day 2 was both reliably above chance but below ceiling in the majority of participants.

#### Episodic encoding session procedure

On the first day, participants completed the episodic encoding phase. This phase presented eight episodes each composed of five unique sequential picture components. Episode components were accompanied with a text segment of a story to encourage the maintenance of the true episode order in memory. The story was written in first-person perspective to better align with veridical personal episodic memories. The first four elements of each episode were taken from 6 potential categories of stimuli: faces, buildings, body parts, objects, animals, and cars. The final element in each episode was not taken from these categories; instead, it represented a unique ending element. Participants were instructed to try to remember the order of the episodes and informed that a bonus would be tied to their performance on questions which tested their memory for the sequential order of the episode elements. A practice episode was presented in the first instance, after which participants were asked to type in the name of the 1^st^ stimulus element presented in an episode, then the 2^nd^, 3^rd^, 4^th^, and 5^th^ elements.

In each episode, participants were presented with the initial picture element along with a segment of story text shown below (**Fig. 1a**; **Table S1**). A grey screen background was used for all experimental phases. The stimulus faded in over 0.5 sec and was then presented with the story text for 2 sec. The text then disappeared and for the remaining 2.5 sec, participants performed a target detection task, pressing the ‘1’ key whenever they saw a small grey square appear at any location over the stimulus (mean of 1 target per stimulus). The stimulus then faded out for 0.5 sec. Total stimulus duration, including fade-in and fade-out, was ∼ 5.5 sec. A grey ‘bokeh’ image faded in as the stimulus faded out. After the stimulus disappeared, participants responded with the ‘up arrow’ key to a series of 1-3 arrow indicators (‘^ ^ ^’) in order to progress to the next element of the episode. If participants did not respond to an arrow within 6 sec, a warning appeared instructing the participant to respond faster. The mean inter-stimulus interval was 6.5 sec (1 sec for short duration episodes; 12 sec for long duration episodes). For the final component of the episode, a white square initially occluded the stimulus and participants then pressed the ‘space’ key to reveal the stimulus and associated story text. After the final component of the episode, a delay of 2 sec was followed by text “Positive ending: you won +£1.00!” or “Negative ending: you lost - £0.50!” depending on whether the story ended in a positive or negative way. Participants were then presented with a probe requiring them to type in the name of a particular episode element (selected pseudo-randomly from elements 1-4). A 30 sec rest period followed each episode. After the completion of the 8 episodes, participants were instructed not to rehearse the episodes or to record the episodes in any way.

Episodes were constructed from a pseudo-random combination of category elements in addition to a final component that was not taken from any of these categories. A brief story text connected the sequence of stimuli into a short story (**Table S1**). The stimuli consisted 40 photographs, taken either from the internet or previous studies from our group, encompassing the following categories: human faces (6), buildings (6), body parts (5), objects (5), animals (5), automobiles (5), and eight final component pictures (4 negative and 4 positive). As noted above, half of the episodes were of a longer duration, achieved via manipulating the inter-stimulus-interval (1 sec or 12 sec). The story in half of the episodes ended in a positive element and half ended in a negative element (**Table S1**). The ordering of long versus short and positive versus non-positive episodes was pseudo-randomized in two counterbalanced orders.

After a 5 min break to obviate a potential influence of temporal proximity on performance for the last episodes, participants completed a short cued retrieval phase that tested recall of the order of the elements presented in each episode. The memory test was brief to minimize additional ‘exposure’ to episode stimuli. Following a practice trial (using stimuli from the practice episode), participants completed 8 trials in the “after” condition and then 8 trials in the “before” condition. Each mini-block of 8 trials was preceded by text indicating the current condition. Participants were shown a picture cue and instructed to retrieve the associated episode in order to make a response about the sequential order of the subsequent answer stimulus. In the after condition, participants attempted to remember what came after (at any point) the cue in the same episode (**Fig. 1b**). In the example after condition trial in **Fig. 1b**, the participant is cued with the harmonica from the above episode. The presented answer, the SUV, indeed followed the harmonica in the episode, so if the participant remembered the episode and order, she should respond with ‘Correct.’ If the sunny sky or bear was presented as the answer, the participant should also responds with ‘Correct.’ If the answer was the man or a stimulus from any other episode, the participant should respond with ‘Incorrect.’ Answers were ‘correct’ for any position after the cue, not just immediately after it. In the before condition, participants attempted to remember what came before (at any point) the cue in the same episode. In both conditions, when the answer picture was presented, participants were shown the response options “Correct” and “Incorrect” in text below the picture. Cues in this memory test were only taken from the second state 2 (of 5 total episode states) in the after condition or the fourth state in the before condition. The answer on half of the trials was correct.

On each cued retrieval trial, the cue picture was presented in full opacity for 0.5 sec and then faded to 0 % opacity across the remaining 5 sec of the retrieval period (**Fig. 1**). Then the response picture was presented. The answer text indicated the mapping between key responses and answers, e.g. “Correct (1)” and “Incorrect (2)”; the left and right text locations were randomly selected on each trial. There was no time constraint on the answer period. After an answer was recorded, following a brief 0.1 sec pause, a 2-level confidence scale (“High” and “Low”) was presented, with the left and right location of options randomized. After a 0.1 sec pause, a fixation period of mean 1.5 sec followed (randomly sampled from the values [1.0, 1.5, 2.0]).

#### MEG session procedure

Participants returned for the MEG scan on the following day. After initial setup in the MEG room, participants were reminded of the instructions for the cued memory phase and completed 4 practice questions (based on the practice episode from the previous day). During scanning, the memory response period was time-constrained. This limit was added to encourage participants to retrieve as much information from memory as possible during the cue period and to facilitate later MEG analysis of neural processes underlying successful retrieval. Participants were instructed to retrieve as best as possible the episodes during the presentation of the cue picture, and in this way they could respond faster (and avoid missed responses) when the answer appeared. Participants were again reminded of the performance bonus based on memory accuracy.

As described above for the memory test on the first day, on each cued retrieval trial, the cue picture was presented in full opacity for 0.5 sec and then faded to 0 % opacity across the remaining 5 sec of the retrieval period (**Fig. 1**). The gradual fade of the cue across the retrieval period was designed to avoid any sharp stimulus offset effects which could negatively affect MEG decoding. Then the answer stimulus was displayed. The text indicating the key response, e.g. “Correct (1)” and “Incorrect (2)”, was randomly presented on the left and right of the screen. If a response was not made in this time period, the warning “Please try to respond more quickly!” was presented for 2 sec. The response picture was presented for 1-3 sec with the duration based on the recent rate of missed trials in the past 20 trials. If participants made no response on more than 14 % of recent trials, the response period was increased in duration by 0.25 sec (with a ceiling of 3 sec). If participants made no response on less than 5 % of recent trials, the answer period was decreased in duration by 0.25 sec (with a floor of 1 sec). After the answer period, following a brief 0.1 sec pause, a 2-level confidence scale (“High” and “Low”) was presented, with the left and right location of options randomized. If a response was not made in time, the warning “Please try to respond more quickly!” was presented for 2 sec. After a 0.1 sec pause, a fixation period of mean 1.5 sec followed (randomly sampled from the values [1.0, 1.5, 2.0]).

In each of 5 blocks in the cued retrieval phase, trials of after and before conditions were separated into mini-blocks of 10-12 trials. Each mini-block was preceded by an instruction screen: “Next: What picture came after (before)?” along with the instruction to press the ‘1’ key to continue. At the mid-point of each block, participants were given a 30 sec pause, followed by a reminder of the current condition and an instruction to press the ‘1’ key to continue. Each of the five blocks of cued retrieval included 43 trials and lasted for approximately 8 minutes. Brief rest breaks were inserted between blocks. In the cued retrieval phase, we collected ∼ 27 trials per episode and ∼ 43 trials per state (episode positions 1 to 5) for a total of 215 trials. For one participant, MEG data were lost for the final memory retrieval block; the remaining 172 trials were analyzed. All trials with a cue from state 1 were after condition trials. All trials with a cue from state 5 were before condition trials. Trials with a cue from state 3 were composed of equal numbers of after and before condition trials, while trials with a cue from state 2 and state 4 were a weighted mixture of after and before condition trials.

The presented answer was correct on ∼ 39 % of trials. The remaining 61 % of trials were incorrect: on 52 % of total trials, the incorrect answer came from another episode and on the remaining ∼ 9 % of total trials, a ‘lure’ answer was presented that was from the same episode but in the incorrect direction as the current condition. For example, in an after condition trial where the cue was from state 3, a picture from that episode in state 1 was presented as the answer. Note that on the first day participants are exposed to the complete episode only one single time. During the memory test, participants see all episode elements again, but at this stage they are provided in an order that mixes elements between different episodes, or elements within the same episode that are out of the true order, and only very rarely are pairs of elements presented in the original order. Trials were presented in a pseudo-random order with the constraint that no episode was queried on sequential trials.

The cued retrieval phase was followed by a functional localizer to derive participant-specific sensor patterns that discriminated each of the 6 categories that made up the episodes by repeatedly presenting each of the 32 unique stimuli. The localizer design was based on previous studies (19, 20). In brief, participants were instructed to read a word shown on the screen, pay attention to the picture that followed, and respond if any grey square targets appeared superimposed over the picture. The instructions were followed by 4 practice trials.

In detail, in a localizer trial, participants were presented with a brief name corresponding to one of the pictures, presented in text on the center of the screen for 2 sec. Participants were instructed to imagine the corresponding picture. The text then disappeared and the named picture appeared on the screen for 0.75 sec. During picture presentation, participants performed a target detection task, responding with a ‘1’ button press if the picture contained a small grey square. Targets were rare events, appearing on 15.4 % of trials. A mean 0.75 sec fixation ITI followed (range 0.25 – 1.25) during which responses were still recorded. If performance on the target detection task fell below 70 % correct (across missed responses and false alarms), a warning was presented: “Please improve your detection of the grey squares!” Finally, as in the cued retrieval phase, a mid-block rest of 30 sec was inserted during each block. After each localizer block, participants were shown yellow ‘stars’ on the screen, ranging from 0-4, depending on their target detection accuracy in the preceding block.

The stimulus pictures were presented in a pseudo-random order, with the constraint that no category repeat in subsequent trials. Each picture from a given category was presented an equivalent number of times, with 78 repetitions per picture category. The localizer was presented in 5 blocks, with 94 trials in the first four blocks and 92 trials in the last block for a total of 468 trials.

Following scanning, participants completed a post-experiment questionnaire that assessed memory strategy and potential mental reordering of the episodes, and also asked participants to try to write down a brief version of each story. The re-ordering question asked “Did you change the order of the stories to make your own story order? 1= never, 5=always”. Participants who responded with a 4 or 5 were considered for exclusion, in conjunction with performance on the memory and sequence memory test. We observed a negative correlation in the full group (prior to exclusions) between response to this question and memory performance in the MEG session.

Finally, participants completed a computerized sequence memory test where they attempted to place the stimuli from a given episode in the correct order. In this phase, stimuli from an episode were presented in a random order on the left side of the computer screen. Participants then moved each stimulus from the left side (starting from the top) into one of 5 empty boxes spread from the left to the right across the screen. Stimuli were moved using the left and right arrow keys and the space bar was used to confirm placement. Accuracy was measured as the mean rate of correct replacement across each location across all episodes.

### MEG acquisition

Participants were scanned while sitting upright inside an MEG scanner located at the Wellcome Centre for Human Neuroimaging, at UCL. A whole-head axial gradiometer MEG system (CTF Omega, VSM MedTech) recorded data continuously at 600 samples per second, utilizing 273 channels (2 original channels of the 275 channels are not operational). Three head position indicator coils were used to locate the position of participant’s head in the three-dimensional space with respect to the MEG sensor array. They were placed on the three fiducial points: the nasion and left and right pre-auricular areas. The coils generate a small magnetic field used to localize the head and enable continuous movement tracking. We also used an Eyelink eye-tracking system to monitor participant’s eye movements and blinks. The task was projected onto a screen suspended in front of the participants. The participants responded during the task using a 4-button response pad to provide their answers (Current Designs), responding with self-selected digits to the first and second buttons.

### MEG Pre-processing

MEG data were processed using MATLAB packages SPM12 (Wellcome Trust Centre for Neuroimaging) and FieldTrip. The CTF data were imported using OSL (the OHBA Software Library, from OHBA Analysis Group, OHBA, Oxford, UK) and down-sampled from 600 Hz to 100 Hz (yielding 10 ms per sample) for improved signal to noise ratio and to conserve processing time. Slow drift was removed by applying a first order IIR high-pass filter at 0.5 Hz.

Preprocessing was conducted separately for each block. An initial preprocessing step in OSL identified potential bad channels whose characteristics fell outside the normal distribution of values for all sensors. Then independent component analysis (FastICA, http://research.ics.aalto.fi/ica/fastica) was used to decompose the sensor data for each session into 150 temporally independent components and associated sensor topographies. Artifact components were classified by automated inspection of the combined spatial topography, time course, kurtosis of the time course, and frequency spectrum for all components. For example, eye-blink artifacts exhibited high kurtosis (>20), a repeated pattern in the time course and consistent spatial topographies. Mains interference had extremely low kurtosis and a frequency spectrum dominated by 50 Hz line noise. The maximum number of potential excluded components was set to 20. Artifacts were then rejected by subtracting them out of the data. All subsequent analyses were performed directly on the filtered, cleaned MEG signal, in units of femtotesla.

In the cued retrieval blocks, an 8.5 second epoch was extracted for potential analysis for each trial, encompassing 500 ms preceding cue onset and continuing past the answer response. In the analyses below, we analyzed the first two-thirds of the cued retrieval period. Given the speeded response demands to the response stimulus, the end of the period is likely to involve increasing response preparation that could decrease the ability to detect sequenceness events. We excluded also the initial 160 ms following cue presentation to allow time for early stimulus processing. Thus, our retrieval period analysis window focused on 160 - 3667 ms of the full 5500 ms period. In the localizer blocks, a 4.5 second epoch was extracted for potential analysis for each trial, encompassing 500 ms preceding text onset through the end of the picture presentation period. In both the retrieval and localizer blocks, preceding the analysis steps described below, we excluded time periods within individual channels that exhibited extreme outlier events (determined by values > 7x the mean absolute deviation).

### MEG data decoding and cross-validation

Lasso-regularized logistic regression models were trained for each category. Methods followed those used in previous studies (19, 20). Only the sensors that were not rejected across all scanning sessions in the preprocessing step were used to train the decoding models. A trained model k consisted of a single vector with length of good sensors n consisting of 1 slope coefficient for each of the sensors together with an intercept coefficient. Decoding models were trained on MEG data elicited by direct presentations of the visual stimuli.

For each category we trained one binomial classifier. Positive examples for the classifier were trials on which that category was presented. Negative examples consisted of two kinds of data: trials when another category was presented, and data from the fixation period before the text pre-cue appeared. An equal number of events of null data were included as there were actual events. The null data were included to reduce a potential correlation between different classifiers – enabling all classifiers to report low probabilities simultaneously.

To examine localizer performance we used cross-validation. We computed the number of included trials per category (after exclusion of trials due to MEG artifacts). We then calculated the number of cross-validation folds by subtracting the minimum number of trials included across categories plus one; the number of folds per participant was usually between 15-20. Classifier performance was estimated on the included data and tested on randomly determined left-out data for N folds; performance was then averaged across folds to derive a mean value.

Separately, for classifying memory retrieval data a different classifier was trained. This classifier was trained on all localizer trial data with no cross-validation; cross-validation is not used for across-phase analyses as no inferences are made based on the localizer performance itself. Prediction accuracy was estimated by treating the highest probability output among all classifiers as the predicted category. Sensor distributions of beta estimates are shown in **Fig. S2** and prediction performance of classifiers trained on 200 ms on left-out trials in functional localizer task are shown in **Fig. S3**.

To determine whether the categories used in the experiment were reflective of how these stimuli were actually represented in the MEG data as participants viewed the stimuli, we conducted a supplemental classification analysis that trained a separate classifier for each of the 32 stimuli (4 category-level stimuli for each of 8 episodes). This analysis used cross-validation as described above. Also as above, a second classification analysis did not use cross-validation but instead trained on the full localizer phase and tested on the memory test phase cue-evoked responses. An alternative classification analysis trained each stimulus versus all other stimuli but left out the other members of that stimulus’ category (e.g. training for face1 omitted trials with face2-face6). Note that the classification of individual stimuli was under-powered given the low number of repetitions per stimulus, and our localizer phase was not designed to produce robust single-stimulus classifiers for use on the sequenceness analyses. These results are detailed in **Supplemental Results**.

### Sequenceness measure

The decoding models described above allowed us to measure spontaneous reactivation of task-related representations during memory retrieval. We next defined a ‘sequenceness’ measure in terms of the degree to which these representations were reactivated in a well-defined sequential order (19, 20). Here we utilized an updated general linear model approach (20). This analysis approach is illustrated in **Fig. S2**. Briefly, the method approximates a lagged cross-correlation between category evidence for transitions in a given episode. As such, the method utilizes the full period of analysis in the calculation and produces a single statistic representing the strength of sequenceness across this full period. Discrete sequential events are not identified, though in theory each retrieval period could include numerous events.

First, we applied each of the six category decoding models to the cued retrieval period MEG data. This yielded six timeseries of reactivation probabilities for each trial, each with length N, where N is the number of time samples included in the retrieval period analysis window. Below, we use the term “stimulus” for simplicity to refer to the category-level information.

We then used a linear model to ask whether particular sequences of stimulus activations appeared above chance in these timeseries. For each stimulus *i*, at each possible time lag Δ*t*, we estimated a separate linear model:

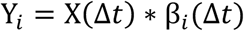

The predictors X(Δ*t*) were time-lagged copies of the six reactivation timeseries. The model predicted Y*_i_*, the reactivation of stimulus *i*. The linear model had N rows, with each row a time sample. We estimated β*_i_*(Δ*t*), a vector of coefficients that described the degree to which stimulus *i*’s reactivation was predicted by activation of each other stimulus at time lag Δ*t*. By repeating this procedure for each stimulus *i*, we obtained β*_i_*(Δ*t*), a 6×6 matrix that can be viewed as an empirical transition matrix between the six stimuli (categories) at lag Δ*t*.

Specifically:

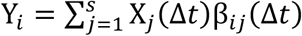

Where X*_j_* (Δ*t*) are time-lagged copies of Y*_j_*, *s* is the number of states, and therefore:

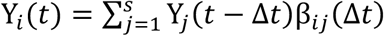

The matrix β*_i_* (Δ*t*) is obtained by solving the following set of equations for each stimulus *i*, up to state *s*.

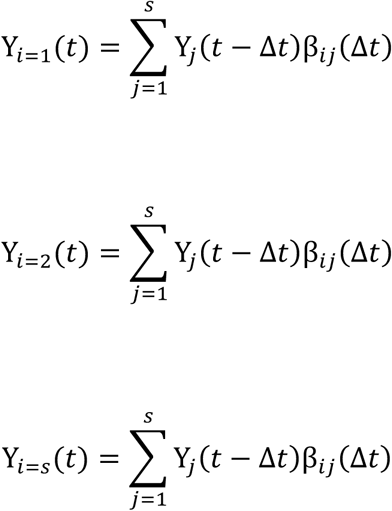

We next asked whether the β*_i_* (Δ*t*) was consistent with a specified 6×6 transition matrix by taking the Frobenius inner product between these two matrices (the sum of element-wise products of the two matrices). This resulted in a single number *Z*_Δ*t*_, which pertained to lag Δ*t*. Finally, differential forward – backward sequenceness was defined as *Z_f_*_Δ*t*_ − *Z_bΔt_*. In our initial analyses and individual differences analyses, we used the difference between correlations in the forward (*Z_f_*_Δ*t*_) and backward (*Z_bΔt_*) direction in order to remove common autocorrelation which would otherwise add significant variance. In the analyses testing for a relationship between sequenceness and trial-by-trial accuracy, we entered the separate forward (*Z_f_*_Δ*t*_) and backward (*Z_bΔt_*) sequenceness measures into the regression analyses. As our analysis was on trial-based data and not rest, we did not need to control for alpha rhythm (20).

The transition matrix was defined as the stimulus (category) order in each episode. Our primary results focus on comparisons of sequenceness on correct versus incorrect retrieval trials; as such, we do not conduct comparisons to a null value. Here, as category orders were pseudo-randomly shuffled across episodes, we did not conduct permutation tests. To ensure that the results were not overfit to the regularization parameter of the logistic regression, all results were obtained with the lasso regularization parameter that yielded the strongest mean decoding in the localizer (l1 = 0.002). The decoding models used to evaluate sequenceness were trained on functional localizer data taken from 200 ms following stimulus onset. The 200 ms time point exhibited the strongest decoding accuracy during the localizer; notably, this time point of category decoding was also consistent with the individual stimulus decoding findings of Kurth-Nelson et al. (19) and Liu et al. (20). We only included trials with a button response to the probe stimulus; all trials with no response were excluded from analysis.

In an initial step, prior to the multilevel modelling analyses, we localized a time lag of interest in the after condition over correct trials using a leave-one-participant-out cross-validation procedure. For a given held-out participant, the absolute value of the peak response across the remaining participants determined the lag for the held-out participant. The analysis included lags from 40-350 ms. These peak times ±10 ms were used to select trial-by-trial sequenceness values.

### Identifying Replay Onsets

Replay onsets were defined as moments when a strong reactivation of a stimulus was followed by a strong reactivation of the next (or preceding) stimulus in the sequence from an episode (20). In this analysis, we first found the stimulus-to-stimulus time lag Δ*t* at which there was maximum evidence for sequenceness (as described above), time shifted the reactivation matrix *X* up to this time lag Δ*t*, obtaining X(Δ*t*). We then multiplied *X* by the transition matrix *P*, corresponding to the unscrambled sequences: *X* × *P*. Next, we element-wise multiplied X(Δ*t*) by *X* × *P*. The resulting matrix had a column for each stimulus, and a row for each time point in the cue period for each trial. We then summed over columns to obtain a long vector *R*, with each element indicating the strength of replay at a given moment in time (across trials). Finally, we thresholded *R* at its 95th percentile to only include high-magnitude putative replay onset events across all trials. We also imposed that constraint that a replay onset event must be preceded by 100 ms of replay-onset-free time.

Specifically:

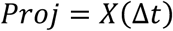

Matrix *Proj* is obtained by time shifting the reactivation matrix *X* to time lag Δ*t*.

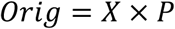

Matrix *Orig* is obtained by matrix multiplication between reactivation matrix *X* and transition matrix *P*.

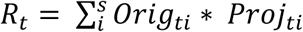

Vector *R* is obtained by elementwise multiplication between matrix *Orig* and *Proj*, and then summing over columns.

### Cue-triggered reactivation analyses

In the cued retrieval period, we tested for cue-triggered reactivation of episode elements. This analysis compared evidence for categories present in a cued episode versus categories not present in a cued episode. The analysis utilized the raw classifier evidence vectors (*n* categories by *t* trial timepoints) to investigate differential activity near the peak stimulus response at ∼ 200 ms. For each episode, the within-episode categories that were not presented as a cue were averaged to derive a measure of reactivation of within-episode elements. In the after condition, there were 3 within-episode categories; in before condition, trials where the cue came from state 5 had 4 categories entered into the within-episode analysis. The 2 categories that were not members of the cued episode were averaged to derive a measure of other-episode reactivation. The timepoints showing the strongest difference between these two measures were averaged for each trial to derive trial-by-trial reactivation measures representing relative within-versus other-element activity. These values were subsequently entered into multilevel regression analyses. We examined a relationship between the trial-by-trial reactivation measure and mean episode accuracy: the average performance across trials for the episode cued on a given trial. We also examined the relationship to trial-by-trial accuracy, but this analysis was under-powered in the very high performing participants. The reactivation analyses collapsed across the after and before conditions.

### Time-frequency analyses

A frequency decomposition (wavelet transformation) was computed for the memory retrieval period in every trial. From this data, we extracted power changes surrounding putative replay onset events.

### Zero-lag correlation analysis

In a supplemental analysis, we examined the relationship between reactivation of within-episode elements compared to other-episode elements with a zero time lag. This measure was a basic correlation between the time series of category evidence: the average of 3 correlations for the within-episode elements and 2 correlations for the other-episode elements. We did not find a greater correlation between within-episode elements than between other-episode elements. Through thresholding of the category evidence time series, we found that correlations were driven by increases in evidence and that these increases were brief (**Fig. S10**). However, we found no relationship between the correlation of within-episode elements across the retrieval period and behavior (**Fig. S10**).

### Multilevel modelling

We conducted all pre-processing of behavioral and MEG data for multilevel modelling in Matlab. Multilevel models were implemented in R, following previous procedures (53). We used a multi-level logistic regression model (glmer, in the lmer4 package) to predict correct memory responses. A correct response in the cued retrieval phase was an answer stimulus correctly identified as coming after the cue in a given episode, an answer stimulus correctly rejected as coming after the cue in a given episode, etc. All missed response trials (where no response was recorded within the response time window) were excluded from analysis.

The primary models included sequenceness derived from the current episode transition matrix. Additional control models examined the effect of sequenceness derived from transitions present in all other episodes but not present in the current episode.

For trial-by-trial accuracy analyses, we included only participants with greater than 10 MEG artifact-free trials in each condition. In general, our exclusion was intended to be conservative and to align with practices in fMRI research regarding approximately sufficient numbers of trials in a condition. We also had a conceptual reason to exclude participants with very few miss trials. In the very high performing participants, miss trials are more likely to be dominated by lapses in attention and resulting error button presses than in regular performing participants; including miss trials in these participants then would add noise to the analyses.

In the main sequenceness analyses, we fit separate intercept, forward sequenceness, and backward sequenceness effects for each participant. In the model, we also included control variables representing performance in neighboring trials. These variables were included because we found that performance 1 and 2 trials in the past and performance 1 and 2 trials in the future was positively related to current trial performance, an effect similar to what we have observed in previous memory studies. In analyses of continuous variables such as mean correct performance for the episode cued on the current trial, we used multi-level regression (lmer).

For all models, to ensure convergence, models were run using the bobyqa optimizer set to 10^6^ iterations. We estimated confidence intervals using the confint.merMod function and p-values using the bootMer function (both from the lmer4 package) using 2500 iterations. All reported p-values are two-tailed.

### MEG Source Reconstruction

All source reconstruction was performed in SPM12 and FieldTrip utilizing OAT. Forward models were generated on the basis of a single shell using superposition of basis functions that approximately corresponded to the plane tangential to the MEG sensor array.

Linearly constrained minimum variance beamforming (54) was used to reconstruct the epoched MEG data to a grid in MNI space, sampled with a grid step of 5 mm. The sensor covariance matrix for beamforming was estimated using data in broadband power across all frequencies.

For the category localizer analysis, the baseline activity was the mean power averaged over 50 ms following stimulus onset. All non-artifactual trials were baseline corrected at source level. We estimated the main effect of each category and contrasts of each category versus all other categories and extracted the peak 200 ms after onset for display.

For the replay onsets analysis, the baseline activity was the mean power averaged over −100 ms to −50 ms relative to replay onset. All non-artifactual trials were baseline corrected at source level. We looked at the main effect of the initialization of replay. This analysis was conducted separately to investigate backward replay events in the after condition and forward replay events in the before condition.

The statistical significance of clusters identified in the beamforming analysis was calculated using SPM12. An initial cluster-forming threshold of p < 0.001 was applied and regions exceeding p < 0.05 whole-brain family-wise-error corrected (FWE) at the cluster level are reported. The timepoint preceding replay onset (−10 ms) was additionally investigated to explore whether individual differences in memory performance related to differential MTL power preceding replay onset.

### Individual differences

We tested for a relationship between MEG measures of sequenceness and mean memory performance in the after and before conditions. For sequenceness, we used differential (forward-backward) sequenceness given the strong decaying autocorrelation evident in the raw forward and backward sequenceness estimates (19, 20). In a supplemental analysis, we estimated the relationship between replay and memory performance using a regression, separately entering forward and backward sequenceness as predictor variables. These analyses used Pearson correlations, reporting two-tailed p-values. A statistical comparison of the correlations between of sequenceness and behavior in the after condition and the before condition was conducted using a test for the difference between two dependent correlations. This test is conservative, as the performance measures in the after condition and the before condition were not identical, while the test assumes full dependence.

We conducted an additional conservative permutation analysis since it is possible that under certain circumstances, having increasing variability in the underlying data toward one end of a distribution across participants might impact on the chance rate of finding a correlation – whether positive or negative – between two variables. Specifically, because lower performing participants have fewer correct trials entered into the mean value than is the case for higher performing participants, the mean values for low performing participants may show higher random variation by chance. (Note that any observed differences in correlation direction between the after and before conditions would not be explained by any such effect.) To conduct the simulation, we pooled all empirical trial-by-trial sequenceness values across participants, separately for the after and before conditions, and mean-corrected the data. From this set of values, we randomly extracted (without replacement) values to match the number of included trials per participant. In each simulated participant, these values were then scaled to match the standard deviation of an actual participant’s trial-by-trial sequenceness data. Across each of 500k simulations, we computed the correlation between mean memory performance and the permuted and scaled sequenceness measure. The resulting p-values were used to determine a conservative permutation-based threshold

### Simulation of MEG analyses and relationship to performance

To provide additional support for our results, we conducted simulations to confirm that the relationship between randomly generated MEG data and behavioral measures is what would expected by chance. All processing and analysis steps were as described above, beyond the generation of simulated MEG data. The simulation proceeded in 3 steps: 1) generation of MEG localizer data and training of classifiers, 2) generation of MEG memory retrieval data, applications of classifiers, and calculation of sequenceness for each trial, and 3) multilevel modelling to relate sequenceness to behavior.

In step 1, we first estimated a sensor covariance matrix based on random data (here and below using the randn function in Matlab), constructed sensor patterns per category, and generated category training data for each category based on random data plus the generated sensor patterns. Classifiers (one per each of 6 categories) were trained on these data.

In step 2, for each trial, MEG data were generated across all sensors using the mvrnd function in Matlab. Across time, an estimated temporal auto-correlation derived from the actual data (0.65) was applied, as well as the previously derived covariance across sensors. Then the sequenceness analysis was applied per trial as in the main analysis described above. This produced a sequenceness measure in the forward and backward direction for each lag up to 350 ms.

In step 3, the values for the simulated after condition on simulated correct trials were extracted for each participant. We then applied a leave-one-out cross-validation procedure for time lag selection. As in the analysis of real data, the lag selected for the left-out participant was based on the peak absolute magnitude of forward minus backward (or differential) sequenceness at lags from 40-350 ms. Across all trials, the mean sequenceness in the forward and backward directions at this peak ±10 ms were entered into the multilevel logistic regression analyses.

One set of simulations utilized all potential behavioral variables from the actual counterbalancing assignment and data (accuracy per trial, exclusion / exclusion of MEG data per trial, after/before condition, and cued episode transition matrix). A second set of simulations approximated the behavioral variables (similar distribution of mean performance across simulated participant, equal number of excluded MEG trials, and equal sampling of each of the episodes). The two simulations based on real behavioral data and simulated behavioral data were each run 10000 times.

### Data availability

Complete behavioral data will be publicly available on the Open Science Framework (https://osf.io/qaewv/). Unthresholded group beamforming statistical parametric maps of replay onset power changes and category responses during the localizer can be found on NeuroVault (https://neurovault.org/collections/6088/). The full MEG dataset will be publicly available on openneuro.org.

### Code availability

Code for the sequenceness analysis, as included in the full processing pathway simulation, is available at: https://github.com/gewimmer-neuro/memory-sequences.

## Supplemental Results

### Sequenceness and individual differences in memory performance

The primary analysis of the relationship between sequenceness and individual differences in memory performance utilized the differential sequenceness measure (fwd – bkw sequenceness; **Fig. 2b-c**). This measure provides a summary of the overall evidence for sequenceness and finds that the same sequenceness direction is important as the trial-by-trial analysis of accuracy. However, given the specific relationship between backwards versus forwards sequenceness in the trial-by-trial analysis of accuracy, we verified that the individual difference relationship was also selective. In the after condition, we found that reverse sequenceness was negatively related to average performance (fwd t_(23)_ = 2.265, p = 0.0337; bkw t_(23)_ = −2.9111, p = 0.0081). In the before condition, we found that forward sequenceness was related to average performance (fwd t_(23)_ = −2.2419, p = 0.0354; bkw t_(23)_ = 1.0456, p = 0.3071). Results from both the after and before conditions show stronger sequenceness in lower-performing participants (reverse sequenceness in the after condition; forward sequenceness in the before condition). These analyses give qualitatively similar results as those reported in the main analysis (**Fig. 2b-c**) which used differential forward-backward sequenceness.

### Sequenceness for ‘other episode’ transitions and trial-by-trial performance

For other episode transitions (including transitions found across the other 7 episodes but not in the currently cued episode), we found that the peak response in correct trials in the after condition using the leave-one-participant-out cross-validation procedure was between 40 and 50 ms (23 participants at 40 ms; 2 participants at 50 ms). We thus examined other episode sequenceness from 40-50 ms. In the model including the other episode sequenceness measure derived from a 40-50 ms lag, we found no significant effects for the other episode measure while the main sequenceness measure remained significant (**Fig. S6**, **Table S4**). We also examined a model including the other episode sequenceness measure from 100-120 ms as a comparison to the main sequenceness lag. Here we also found no significant effects for the other transition measure while the main sequenceness measure remained significant (**Fig. S6**, **Table S5**). Finally, in a separate model looking only at current episode sequenceness at the 40-50 ms lag identified in previous studies and for the other episode measure, we find no relationship between sequenceness and retrieval success in the after condition (p-values > 0.17) or before condition (p-values > 0.52).

### Analyses using single-stimulus classification

As expected, performance of stimulus-level classification during the localizer phase for the cross-validated analysis was markedly lower than performance for the category-level classification during the localizer phase (**Fig. S3**). We also examined cross-classification to the memory retrieval phase (**Fig. S3**). While performance was above the pre-trial baseline level and significant (t_(24)_ = 9.20, p<1e-8), in comparison to the category-level cross-classification results in **Fig. 1f**, the magnitude of the effect versus baseline is much weaker: the effect for the category across-phase classification of memory cues was significantly stronger than the single-stimulus across-phase classification (t_(24)_ = 12.47, p<1e-11). This relatively poor performance when category was ignored during training was expected, given that category information is likely to account for the most variance in stimulus responses.

Even though the classifier showed cross-classification performance that was numerically close to chance, we nevertheless examined whether a sequenceness measure derived from single-stimulus classification might show a relationship to memory retrieval success. In the after condition, we found no relationship between single-stimulus backward sequenceness and retrieval success (p = 0.447). However, in the before condition we found a positive, but non-significant, relationship between single-stimulus forward sequenceness and retrieval success (p = 0.0913).

## Supplementary Figures and Tables

**Figure S1.**
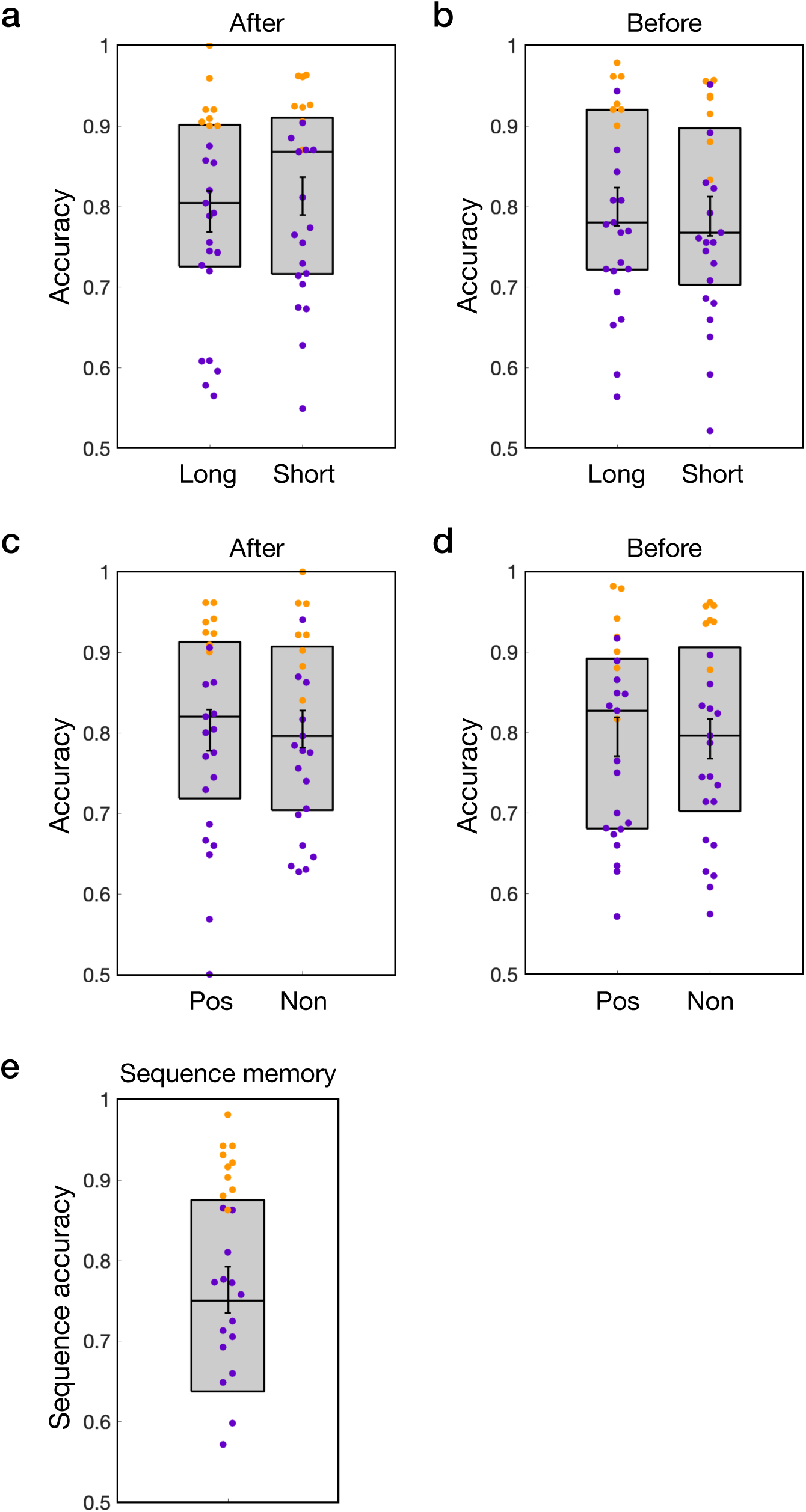
Memory performance as a function of episode length and whether the episode ended in a positive or negative element and performance on final episode re-ordering test. As in **Fig. 1c**, the data points for regular performance participants are represented in purple and very high performance participants are represented in orange. (**a** and **b**) Memory did not significantly differ in the after condition by length (t_(24)_ = −1.389; p = 0.178; TOST equivalence test p = p = 0.065, thus we are unable to rule out the presence of a medium-sized effect) or the before condition by length (t_(24)_ = 0.661; p = 0.515; TOST equivalence test p = 0.0156). (**c** and **d**) Memory did not differ in the after condition by end valence (t_(24)_ = −0.068; p = 0.946; TOST equivalence test p = 0.004) or the before condition by end valence (t_(24)_ = 0.1478; p=0.88; TOST equivalence test p = 0.005). Given the null behavioral differences, primary MEG analysis collapsed across these variables. (**e**) Performance on the post-scan episode sequence memory re-ordering test (n=24 participants with sequence test data). Individual scores were the average of accurate placements of each element within each episode. Sequence memory did not have a condition, so regular performance participants (purple) represent those participants included in both the after and before condition regular performance groups (n = 15); the data points for the remaining high performance participants are depicted in orange. (Error bars represent SEM.)

**Figure S2.**
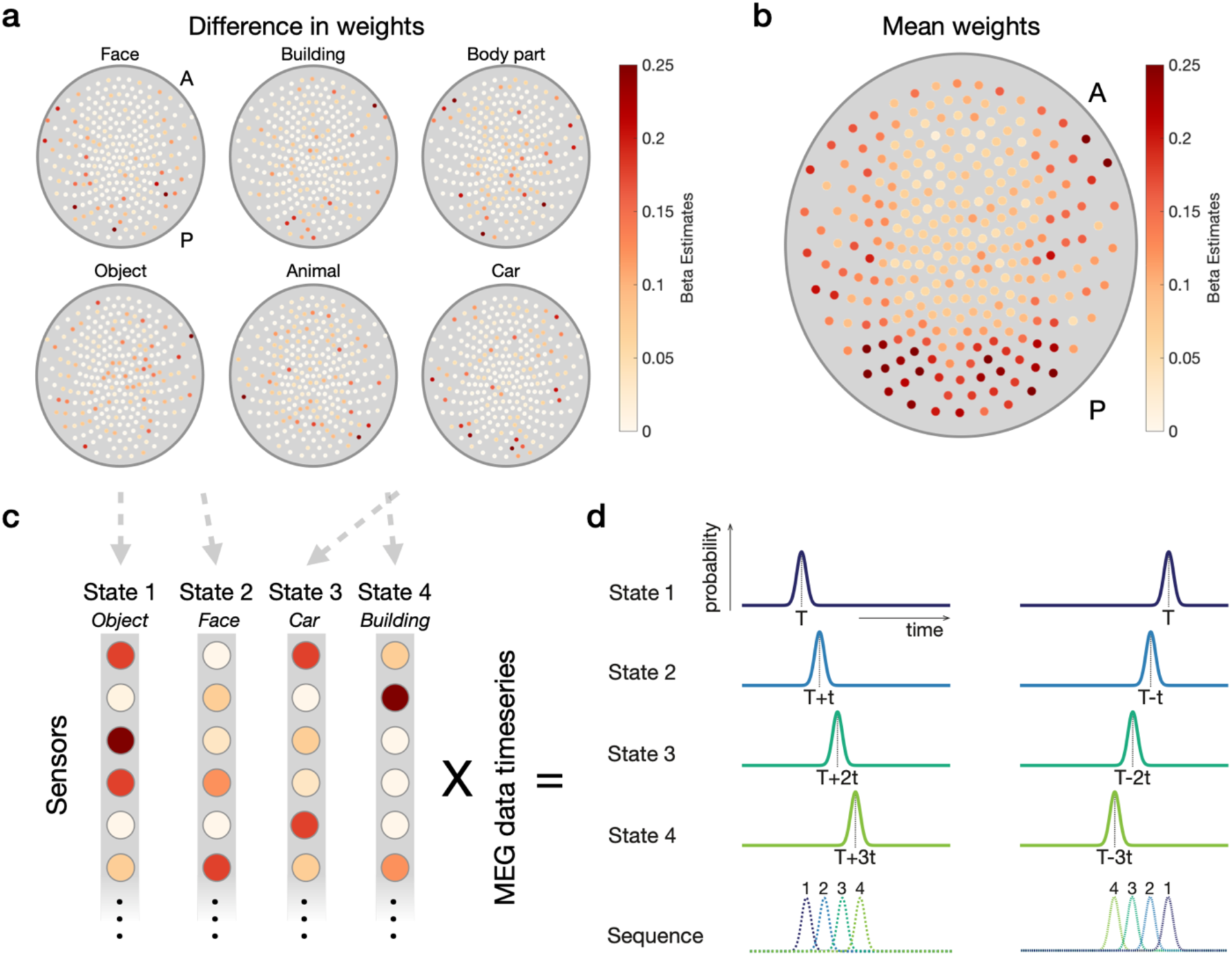
Sequenceness analysis schematic and classifier sensor weighting. (**a**) Classifiers were trained on the 6 categories that made up the episodes. The mean weighting (approximate importance) of each sensor for a given category, minus the mean across all other categories, for illustration only. (Anterior = top; posterior = bottom.) (**b**). Mean sensor weighting across all categories. (**c**) Illustration of how the trained classifiers are applied to the MEG data timeseries for each cued retrieval period, where state 1 - 4 represents episode components 1-4 from **Fig. 1a**. (**d**) The sequenceness analysis detects systematic time shifts (T) in category evidence. A forward sequence illustration is shown on the left; a backward sequence illustration is shown on the right.

**Figure S3.**
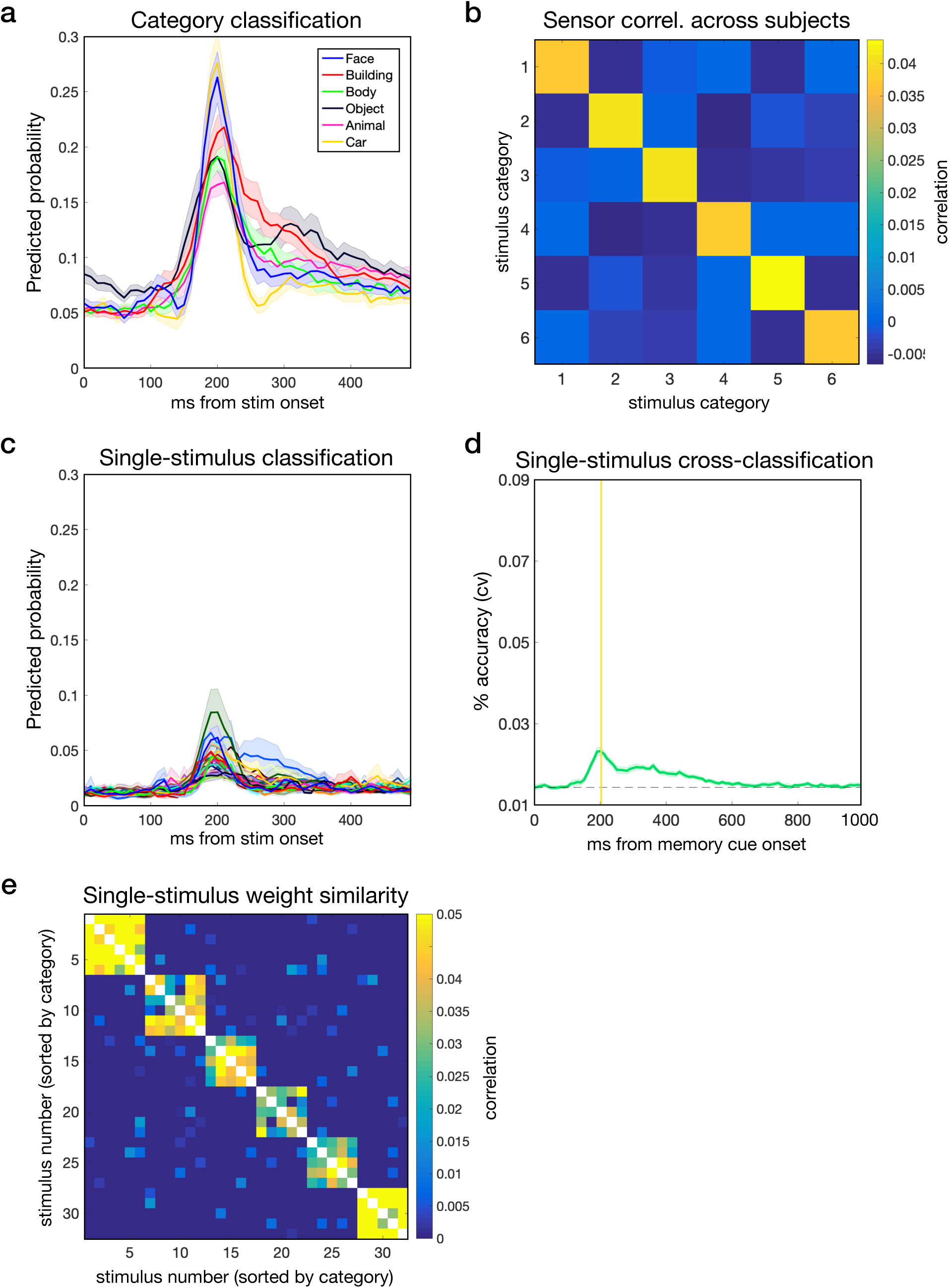
Illustration of localizer classifier performance for the six stimulus categories that made up the first 4 components of episodes (face, building, body part, object, animal, and car) and performance of a classifier trained for each of the 32 individual stimuli from these categories. (**a**) Cross-validated classification performance for each category. Results represent training on the 200 ms time point and testing across all time points. (**b**) Classifier sensor weight correlations across participants within and between-categories reveal strong within-category similarity, suggesting similar sensor importance across participants for the same categories. Categories are sorted as in the legend for panel a: face, building, body part, object, animal, and car. (Lasso regularization parameter set to 1e-6 to maximize sensor inclusion.) (**c**) Classification performance for each of 32 stimuli. Results represent training on the 200 ms time point and testing across all time points. (Colors were randomly assigned.) (**d**) Cross-classification performance for 32 individual stimuli where the classifier was trained on the localizer phase (at 200 ms) and tested on the cues in the memory phase. Compare to the category level cross-classification in **Fig 1f** (y-axis range is matched across figures for comparison). The dashed line represents the maximum classifier value during pre-trial baseline; in statistical tests performance was compared to this baseline value. (**e**) Trained classifier beta weight correlation across sensors across all 32 individual stimuli depict natural emergence of category structure. The image represents that average of individual participant correlation matrices. (Lasso regularization parameter set to 1e-6 to maximize sensor inclusion.) Categories are sorted as in the legend for panel a: face, building, body part, object, animal, and car. (Shaded error margins represent SEM.)

**Figure S4.**
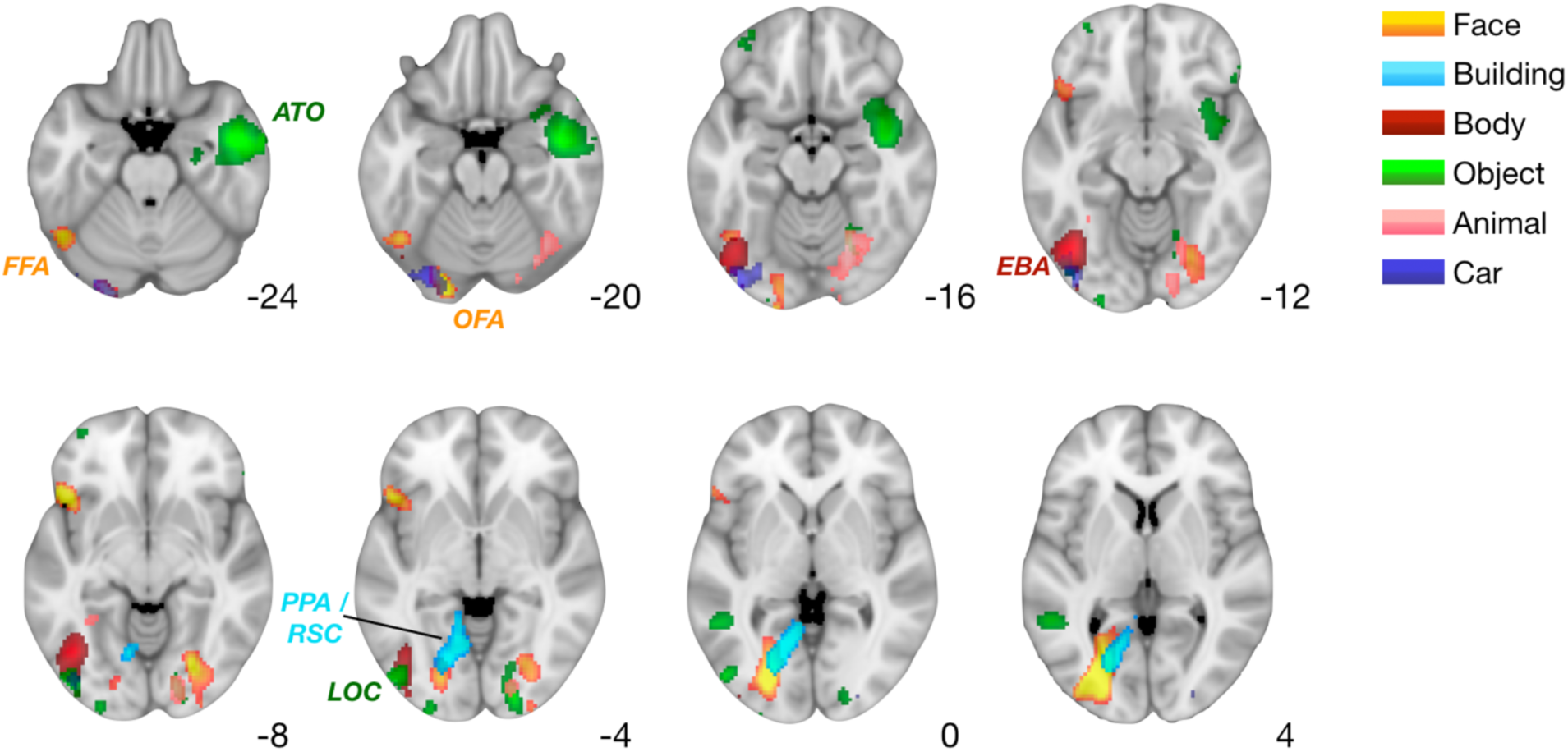
Source localization results for the six categories of stimuli in the localizer phase. Below, each category was contrasted versus all other categories. We found expected patterns of activation for the 4 categories that have received the most investigation in the literature: faces, buildings, body parts, and objects. For faces, activation peaked in a region roughly consistent with the fusiform face area (FFA) as well as the occipital face area (OFA). Activation for building stimuli was located between the well-known parahippocampal place area (PPA) and the retrosplenial cortex (RSC), a region also known to respond to scene and building stimuli. Activation for body part stimuli was in a region consistent with the extrastriate body area (EBA). Activation for objects was in a region consistent with the object-associated lateral occipital cortex (LOC) as well as an anterior temporal cluster that may relate to conceptual processing of objects. Activity for the two less-studied categories, animals and cars, was localized to different areas of the ventral and posterior occipital cortex. Individual category maps thresholded to display localized peaks for illustration. Full unthresholded maps can be found at https://neurovault.org/collections/6088/.

**Figure S5.**
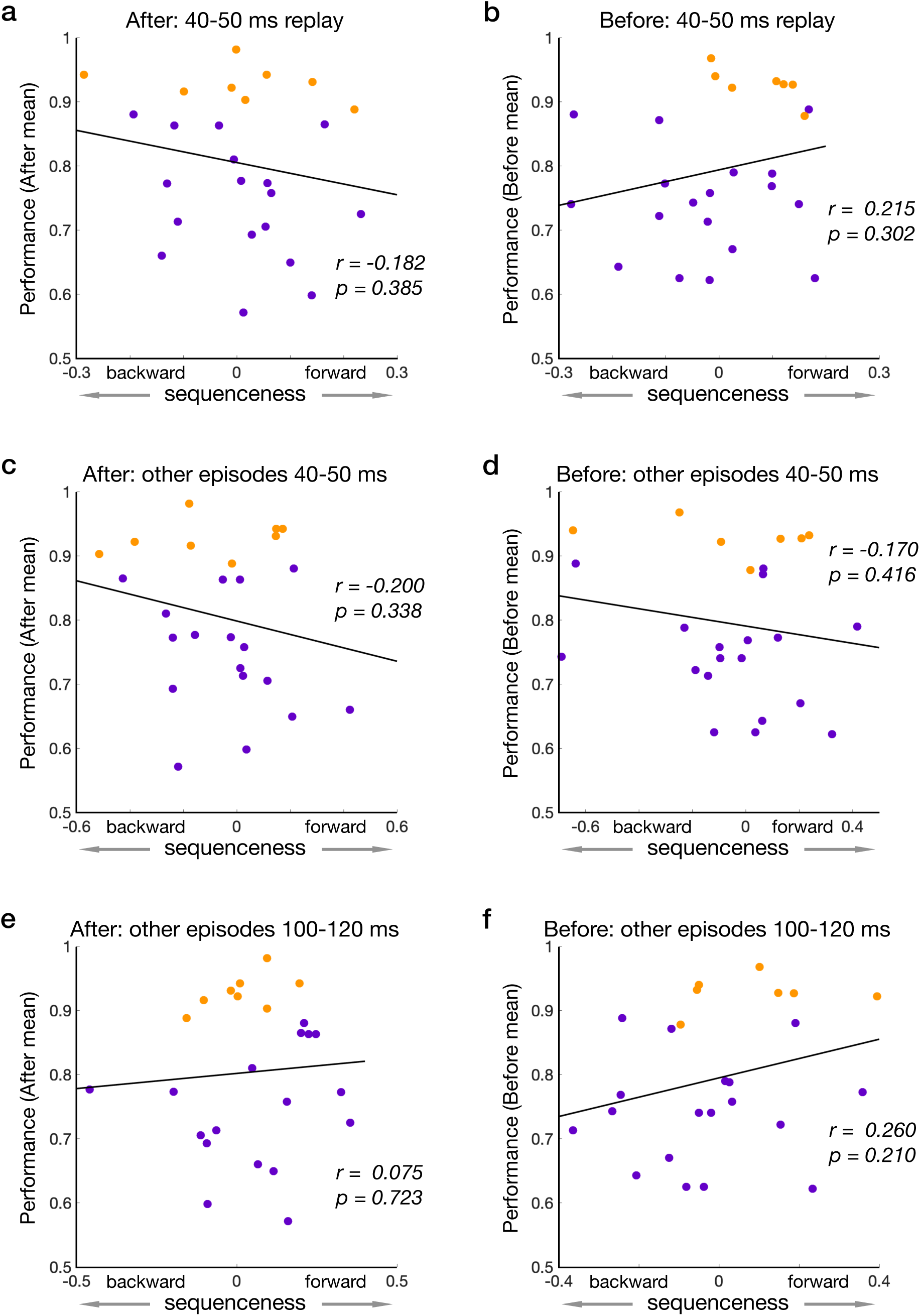
No significant relationship between sequenceness and trial-by-trial behavior at other time lags and in the analysis testing for sequences present in other (non-cued) episodes. (**a-b**) In the after and before conditions, mean sequenceness strength (forward-backward) with a 40-50 ms lag did not relate to overall mean memory performance (percentage of correct trials). As in **Fig. 1c** the data points for the regular performance participants are shown in purple; high performance participants are shown in orange. (**c**) As in panel a, here for the before condition. (**c-d**) In the after and before conditions, mean 40-50 ms sequenceness for other episode transitions (excluding the current episode) did not relate to mean memory performance. (**e-f**) In the after and before conditions, mean 100-120 ms sequenceness for other episode transitions did not relate to mean memory performance.

**Figure S6.**
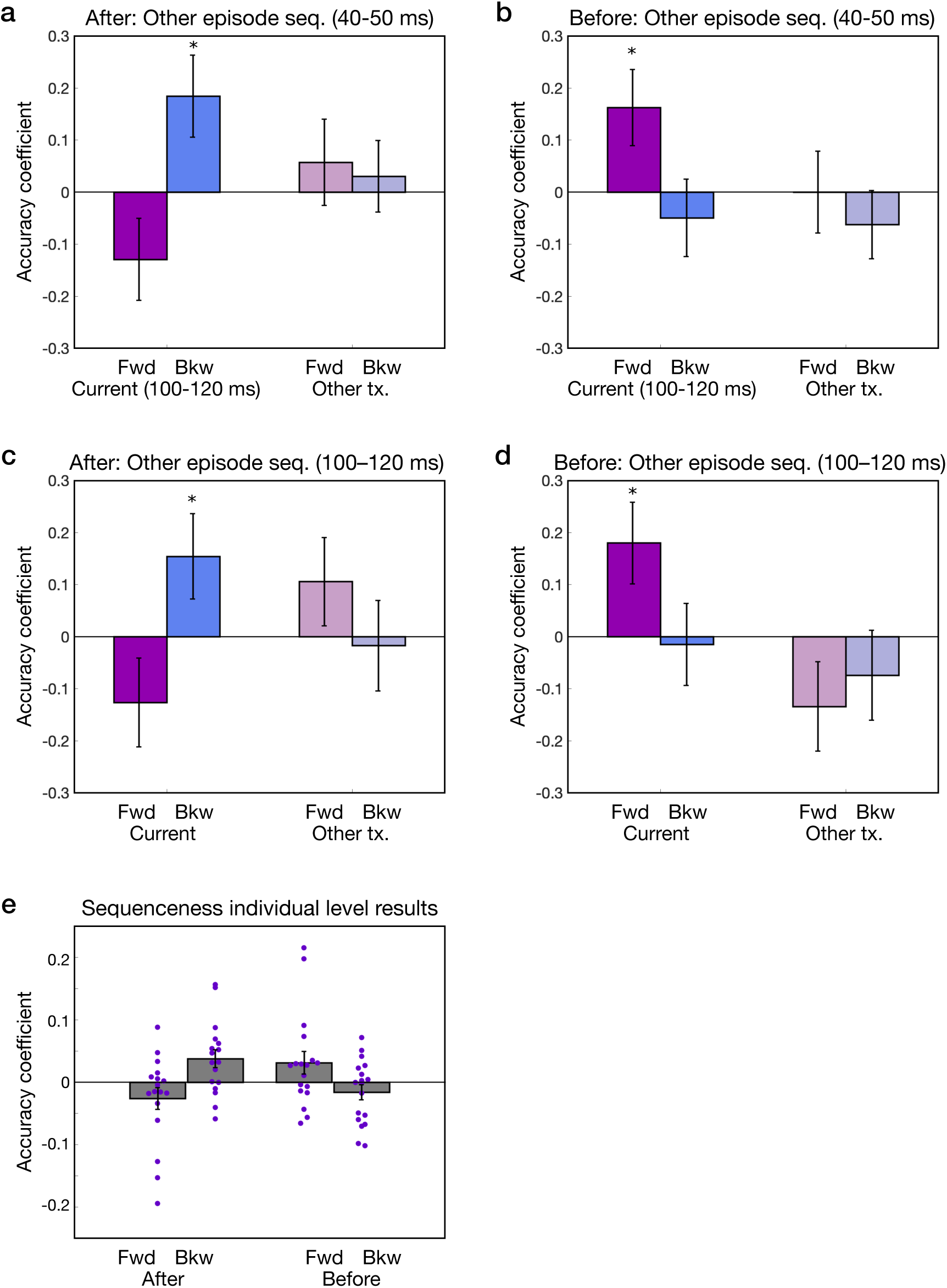
Analyses relating both current episode sequenceness and ‘other episode’ sequenceness to accuracy; individual participant regression results. (**a**) In the after condition, current episode sequenceness (100-120 ms; left, darker color) remains significant (left) while other episode sequenceness at a lag of 40-50 ms shows no relationship to successful retrieval (right, lighter color; **Table S5**). (**b**) As in panel a, but for the before condition. (**c**) In the after condition, sequenceness derived from current episode sequenceness (100-120 ms; left) remains significant while other episode sequenceness (derived from all other transitions excluding the current episode transitions) at a lag of 100-120 ms shows no relationship to successful retrieval (right; **Table S5**). (**d**) As in panel c, but for the before condition. (**e**) Individual regression coefficients for the trial-by-trial relationship between sequenceness and successful retrieval in the after and before conditions as in **Fig. 3a**., but derived from a single-level GLM.

**Figure S7.**
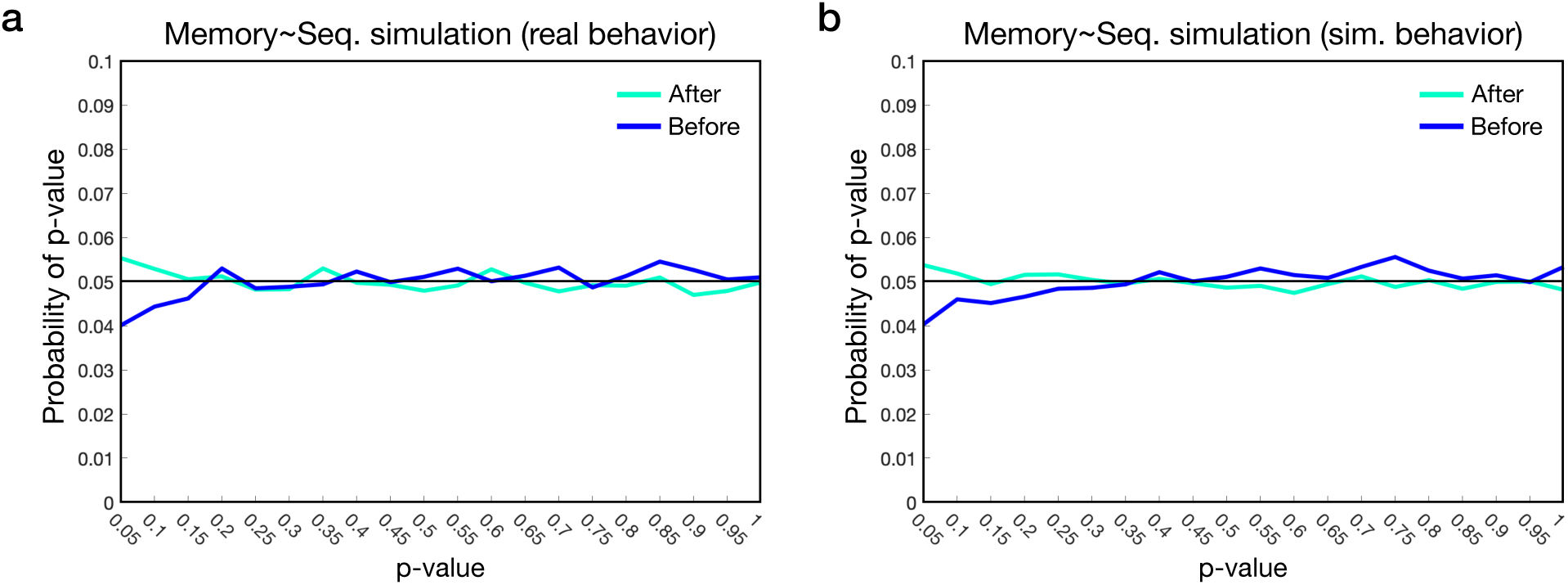
Results of simulating the complete MEG processing and analysis pipeline, showing the relationship between sequenceness and trial-by-trial retrieval success. The panels show the resulting distribution of p-values derived from the sequenceness-retrieval success multilevel regression model across 10k simulations. Simulated p-values were near the 5 % level using both real or simulated behavioral data. (**a**) In the simulations with behavioral variables taken from actual participant data, p-values were equal to or less than 5% in the *after* condition at a rate of 0.055 and in the *before* condition at a rate of 0.04. (After condition in cyan; before condition in blue) (**b**) In the simulations with simulated behavioral variables, the simulated p-value was equal to or less than 5% in the *after* condition at a rate of 0.054 and in the *before* condition at a rate of 0.040.

**Figure S8.**
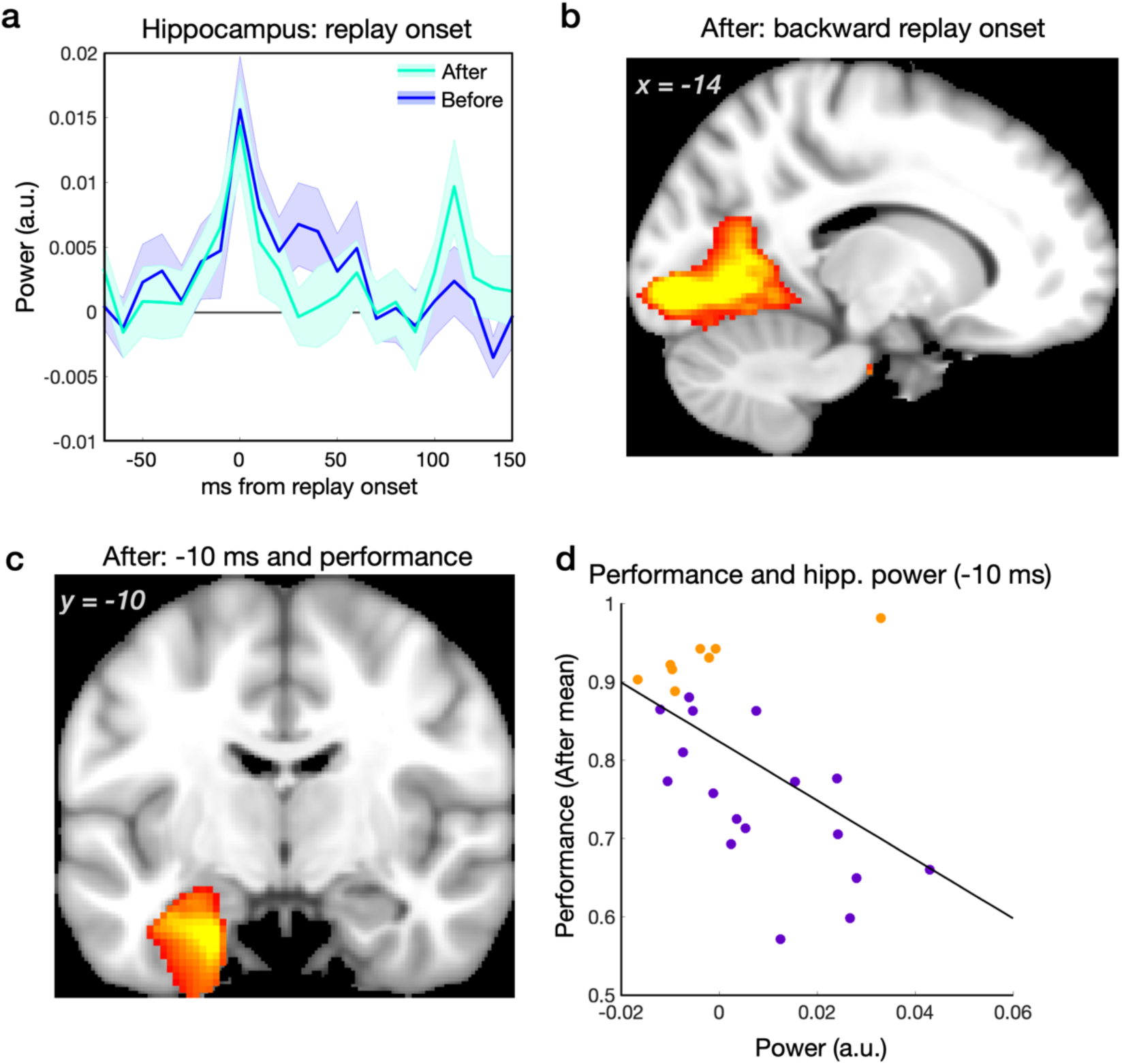
Additional replay onset beamforming results. (**a**) Timecourse of power changes relative to replay onset in the anterior hippocampus in the after (cyan) and before (blue) conditions. (Shaded error margins represent SEM.) (**b**) Power in the right visual cortex at replay onset in the after condition, displaying a different view of the whole-brain results shown in a coronal section in **Fig. 4a**. (Statistical maps thresholded at p < 0.001 uncorrected, for display.) (**c**) Power in the left MTL 10 ms before the onset of reverse sequenceness events correlated with performance, such that lower performing participants showed the strongest increase in power (https://neurovault.org/images/306232/). (**d**) Illustration of performance – power relationship in the right anterior hippocampus. Data are for visualization purposes only and represent the peak coordinate as in panel c. High performance participants in orange; regular performance participants in purple.

**Figure S9.**
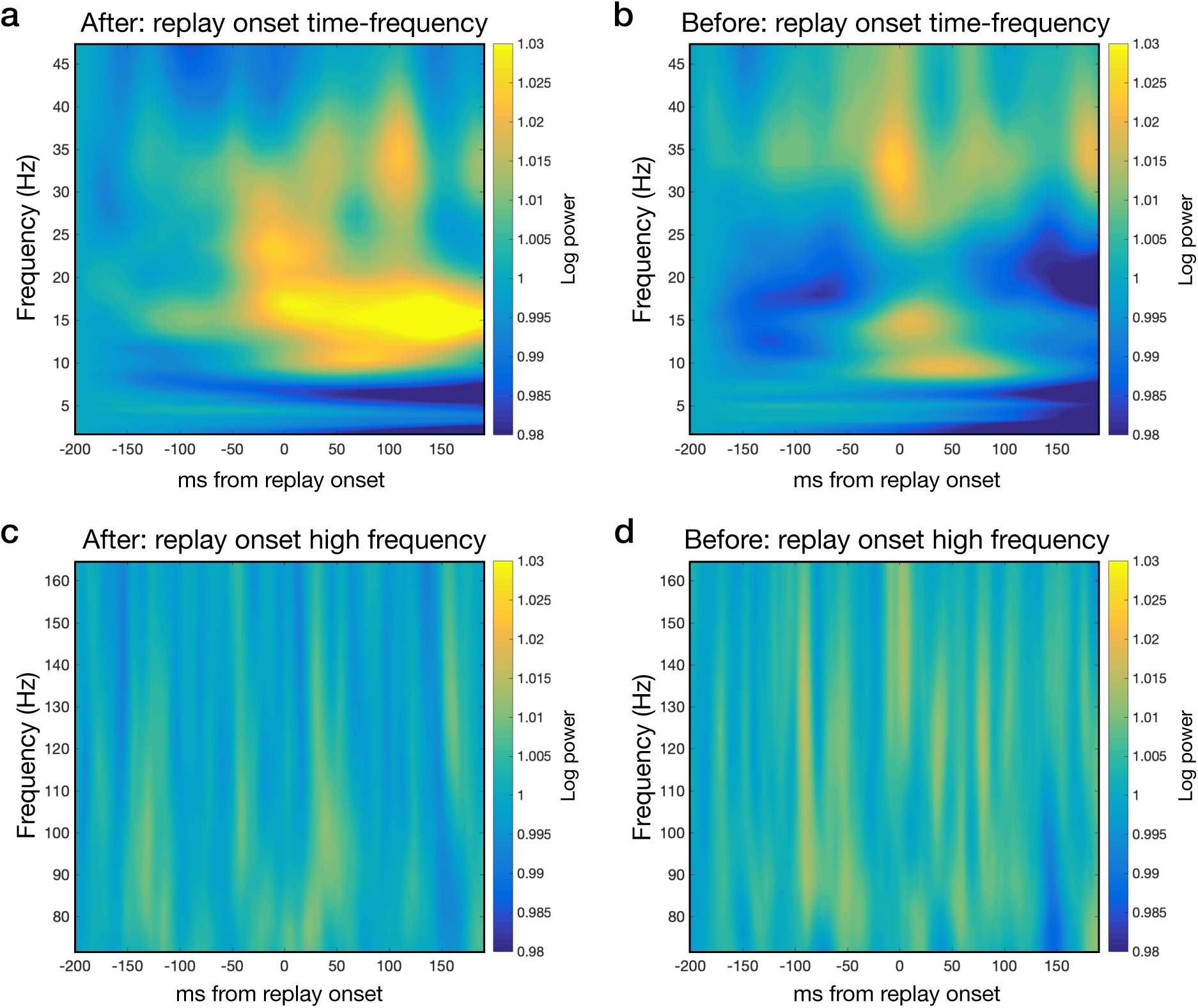
Time-frequency analysis of replay onsets in the after and before conditions separately (**a**) Time-frequency analysis showing power increases at replay onset in the after condition showing frequencies up to ∼ 50 Hz. 0 ms represents the onset of putative replay events. (Average across all n=25 participants in correct trials.) (**b**) Time-frequency analysis as in panel a, here in the before condition. (**c**) Time-frequency analysis of high frequencies in the after condition (using data sampled at 600 Hz) relative to replay onset (**d**) Time-frequency analysis of high frequencies as in panel c, here in the before condition.

**Figure S10.**
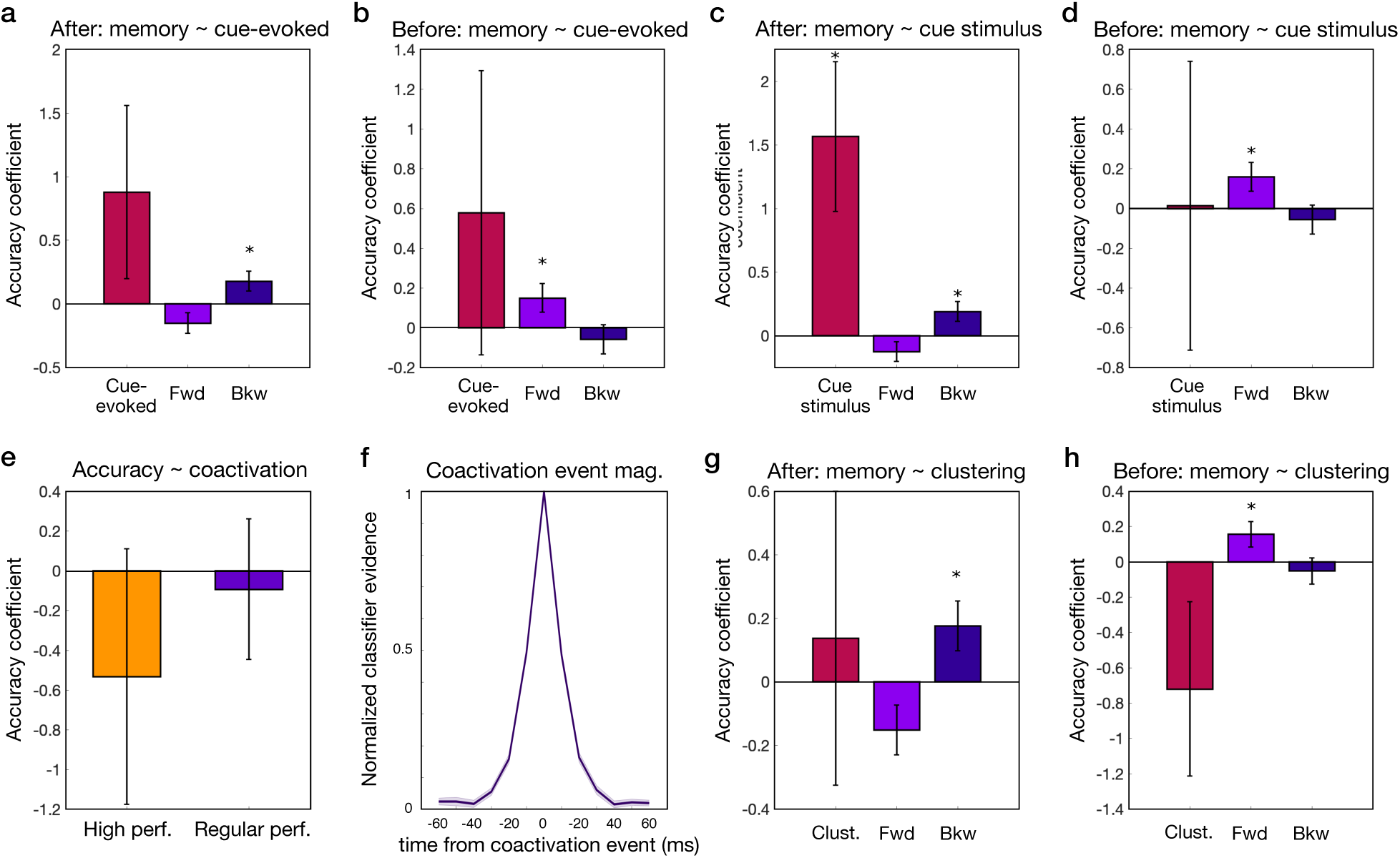
Relationship between accuracy, cue-evoked reactivation, cue response, and zero-lag correlation between within-episode category evidence during the retrieval period. (**a-b**) Cue-evoked reactivation of within-episode elements minus other-episode elements from 200-250 ms and retrieval success in the after condition (**a**) and before condition (**b**), included in the regression model with forward and backward sequenceness. The effects of cue-evoked reactivation were non-significant (**Table S6**); the relationships between sequenceness and memory were unaffected. (**c-d**) Cue response from 200-250 ms and retrieval success in the after condition (**c**) and before condition (**d**), included in the regression model with forward and backward sequenceness. The effect of cue response was significant in the after condition but not the before condition (**Table S6**); the relationships between sequenceness and memory were unaffected. (**e-f**) The correlation between evidence for within-episode categories minus the correlation between all other pairings (zero-lag correlation) across the 160 ms – 3667 ms cue period of analysis (**e**) is not related to trial-to-trial accuracy in very high or regular performance participants: High performance (−0.534 ± 0.644; z = −0.829, p = 0.407); regular performance (−0.093 ± 0.354; z = −0.263, p = 0.792). (**f**) The correlation between within-episode category evidence is driven by high-magnitude events (>= 95 % of mean), and activity for these events peaks and falls rapidly. The purple line represents the mean across participants in the after condition. (**g-h**) The zero-lag correlation between evidence for within-episode categories minus the correlation between all other pairings included in the regression model with forward and backward sequenceness in the after condition (**g**) and in the before condition (**h**). The effect of clustered reactivation was non-significant (after: 0.137 ± 0.463; z = 0.296, p = 0.767; before: −0.721 ± 0.494; z = −1.460, p = 0.144); the relationships between sequenceness and memory were unaffected. (Error bars represent SEM.)

**Table S1.**
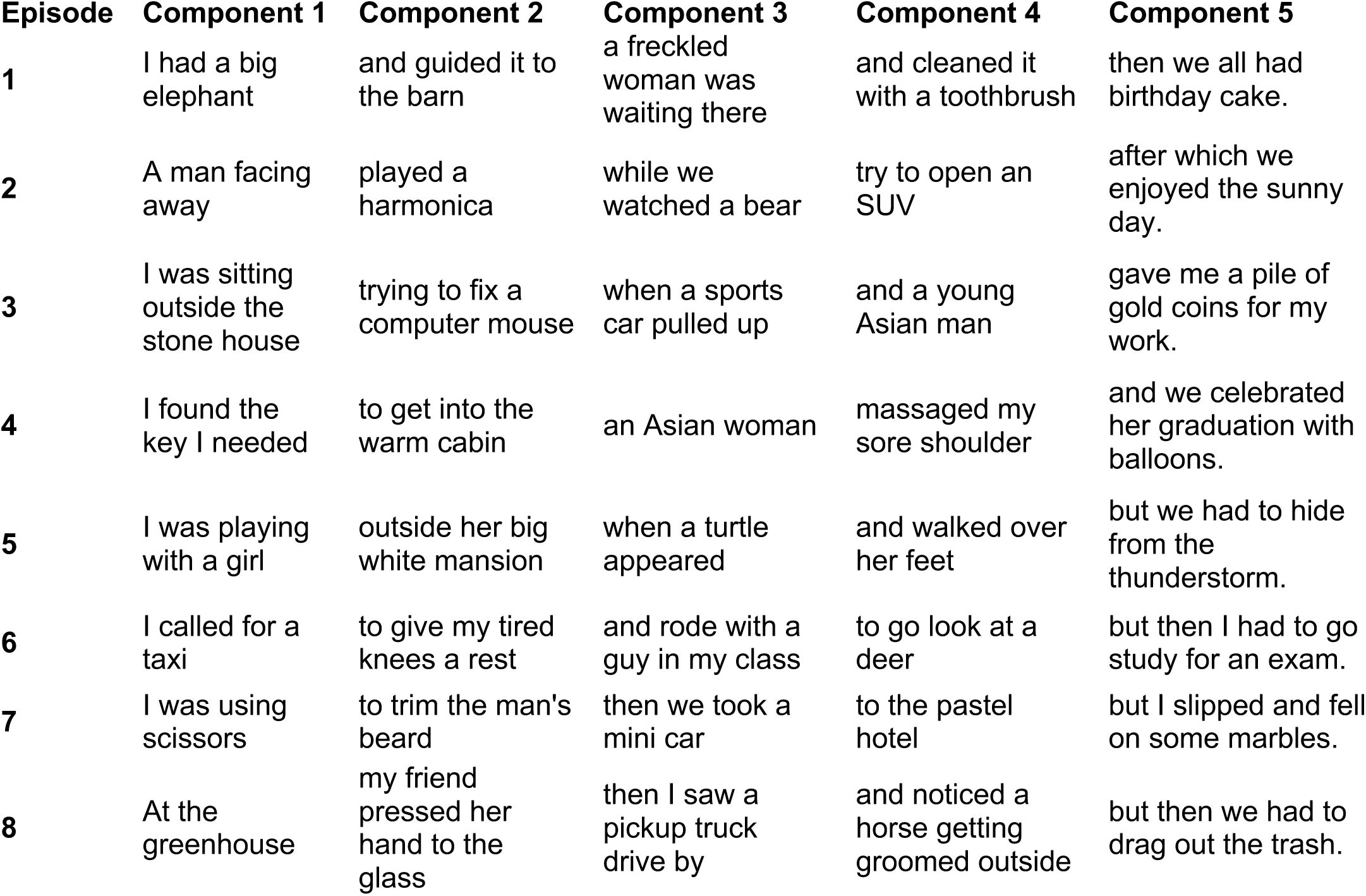
Story text example used in the episodic memory encoding phase on the first day. The stimuli for the first 4 components were taken from the categories: face, building, body party, object, animal, and car. The alternative counterbalance order changed component 5 across episodes from positive to negative.

**Table S2.** Post-experiment written questionnaire answers to questions about memory retrieval. The second column represents the free-form answers to the question “How were you able to remember the associations for the memory questions?”. The third column, labeled ‘Static’, represents the answers to the question “Today, did your memories appear in mind as a sequence or story through time, or did the pictures all appear together as a single combined “static” memory? Single(static) = 1, Sequence (story)=5”. The fourth column, labeled ‘Cue’, represents the answer to the question “When remembering, did you mostly use the period during the initial (fading) picture “cue” to remember, or more the time when the answer option appeared? During the cue = 1, at the answer=5”.

| Num | How were you able to remember the associations for the memory questions? | Q: Static | Q: Cue |
| --- | --- | --- | --- |
| 311 | I tried to quickly "say" to myself the pictures before/after the one on the screen. Then I concentrated whether I see them. | 2 | 2 |
| 312 | Mostly "gut" feeling. | 2 | 2 |
| 313 | Relating some pictures. | 1 | 3 |
| 314 | Some stories that were more relatable to my life were easier to recall. | 5 | 3 |
| 315 | pretty well, it improved over time, but sometimes my concentration slipped, or I pushed wrong button. | 3 | 1 |
| 316 | Trying to recall the story and the order in which the pictures occurred in said episode. | 4 | 1 |
| 317 | I just briefly recalled the stories and quickly ran through the associated images in the episode before seeing the question. | 4 | 2 |
| 318 | Based on the "stories". | 4 | 4 |
| 319 | My own creations/storyline. | - | 1 |
| 320 | Using stories, mnemonics. | 4 | 2 |
| 322 | By remembering the stories or scene which linked the 5 pictures and their order. | 4 | 2 |
| 323 | From the story/associations. | 2 | 4 |
| 324 | Trying to remember the story. | 2 | 1 |
| 325 | I don't think I am able to remember the whole episode, but most of the time, I would remember half. | 4 | 3 |
| 326 | I remember parts of the story line but also remembering which picture was connected to the specific episode. | 4 | 1 |
| 327 | Remember the sequence of events per scenario as a story. | 5 | 1 |
| 328 | I tried to remember the stories and the names you gave to the faces. I only remembered some stories so I guessed the other ones through a process of elimination. | 4 | 4 |
| 329 | I tried to remember the stories with the episode. | 2 | 1 |
| 331 | Using the stories from yesterday. | 5 | 1 |
| 333 | For around half of the stories, it was by eidetic recall. For the other, gut instinct. | 3 | 3 |
| 334 | I tried my best to remember the provided story lines. | 4 | 2 |
| 335 | Based on the stories I made up from the day before. | 5 | 2 |
| 336 | Tried to remember the stories. | 4 | 2 |
| 337 | Own story, verbal encoding e.g. faces looked like some people I knew, etc. | 2 | 3 |
| 338 | Pictured by picture recall and occasionally gut feeling. | 5 | 2 |

**Table S3.** Multilevel modeling results for the inclusion of all participants in the model relating sequenceness and accuracy and effect of task time (trial). Top: model including all participants (with lag selecting using leave-one-out cross validation). Bottom: model including interaction of sequenceness and task time (trial) in regular performing participants.

| **After condition (all $n = 25$)** |  |  |  |  |
| --- | --- | --- | --- | --- |
| **Accuracy ~** | **coef.** | **ste** | **z-stat** | **p-value** |
| *Intercept* | 1.2226 | 0.1594 | 7.671 | <0.0001 |
| SeqFwd | -0.1543 | 0.0718 | -2.148 | 0.0280* |
| SeqBkw | 0.1676 | 0.0713 | 2.35 | 0.0168* |
| **Before condition (all $n = 25$)** |  |  |  |  |
| *Intercept* | 1.374 | 0.1698 | 8.091 | <0.0001 |
| SeqFwd | 0.176 | 0.0685 | 2.571 | 0.0152* |
| SeqBkw | -0.0456 | 0.069 | -0.661 | 0.4912 |
| **After condition: interaction with trial** |  |  |  |  |
| *Intercept* | 0.8586 | 0.1139 | 7.539 | <0.0001 |
| SeqFwd | -0.1403 | 0.0788 | -1.78 | 0.0744 |
| SeqBkw | 0.1705 | 0.0791 | 2.155 | 0.0296* |
| Trial | -0.1274 | 0.1351 | -0.943 | 0.3696 |
| SeqFwd * Trial | -0.0937 | 0.1288 | -0.728 | 0.4704 |
| SeqBkw * Trial | -0.1573 | 0.1337 | -1.176 | 0.2496 |
| **Before condition: interaction with trial** |  |  |  |  |
| *Intercept* | 0.9672 | 0.112 | 8.634 | <0.0001 |
| SeqFwd | 0.1643 | 0.0735 | 2.235 | 0.0248* |
| SeqBkw | -0.0548 | 0.0746 | -0.734 | 0.4232 |
| Trial | 0.1543 | 0.1359 | 1.135 | 0.2544 |
| SeqFwd * Trial | -0.0885 | 0.1197 | -0.739 | 0.4408 |
| SeqBkw * Trial | -0.1147 | 0.1209 | -0.949 | 0.3544 |

**Table S4.** Multilevel modeling results for the relationship between sequenceness from all transitions found in ‘other’ episodes (40-50 ms; excluding transitions from the current episode) to accuracy. Bottom: the preceding model plus the primary measure of sequenceness from current episode transitions (100-120 ms).

| **After condition:** Other transition measure (40-50 ms) |  |  |  |  |
| --- | --- | --- | --- | --- |
| **Accuracy ~** | **coef.** | **ste** | **z-stat** | **p-value** |
| *Intercept* | 0.0617 | 0.08417 | 7.696 | <0.0001 |
| SeqFwd ‘Other’ Tx | 0.06168 | 0.08417 | 0.733 | 0.4592 |
| SeqBkw ‘Other’ Tx | 0.03377 | 0.06787 | 0.498 | 0.6176 |
| **Before condition:** Other transition measure (40-50 ms) |  |  |  |  |
| *Intercept* | 0.9713 | 0.11101 | 8.749 | <0.0001 |
| SeqFwd ‘Other’ Tx | 0.00049 | 0.07813 | 0.006 | 0.9792 |
| SeqBkw ‘Other’ Tx | -0.05764 | 0.06532 | 0.882 | 0.3856 |
| **After condition:** Current + other transition (40-50 ms) |  |  |  |  |
| *Intercept* | 0.86205 | 0.11319 | 7.616 | <0.0001 |
| SeqFwd | -0.12962 | 0.07888 | -1.643 | 0.1024 |
| SeqBkw | 0.18399 | 0.07862 | 2.340 | 0.0240* |
| SeqFwd ‘Other’ Tx | 0.05676 | 0.08312 | 0.683 | 0.5024 |
| SeqBkw ‘Other’ Tx | 0.03009 | 0.06877 | 0.438 | 0.6848 |
| **Before condition:** Current + other transition (40-50 ms) |  |  |  |  |
| *Intercept* | 0.97577 | 0.11245 | 8.677 | <0.0001 |
| SeqFwd | 0.16231 | 0.07315 | 2.219 | 0.0360* |
| SeqBkw | -0.04968 | 0.07465 | -0.666 | 0.4968 |
| SeqFwd ‘Other’ Tx | -0.00061 | 0.07878 | -0.008 | 0.9936 |
| SeqBkw ‘Other’ Tx | -0.06276 | 0.06570 | -0.955 | 0.3528 |

**Table S5.** Multilevel modeling results relating the effect of sequenceness from all transitions found in ‘other’ episodes (100-120 ms; excluding transitions from the current episode) to successful retrieval. Bottom: the preceding model plus the primary measure of sequenceness from current episode transitions (100-120 ms).

| **After condition:** other transition measure (100-120 ms) |  |  |  |  |
| --- | --- | --- | --- | --- |
| **Accuracy ~** | **coef.** | **ste** | **z-stat** | **p-value** |
| *Intercept* | 0.8522 | 0.1112 | 7.662 | <0.0001 |
| SeqFwd ‘Other’ Tx | 0.1414 | 0.0791 | 1.788 | 0.0832 |
| SeqBkw ‘Other’ Tx | -0.0527 | 0.0787 | -0.669 | 0.4968 |
| **Before condition:** other transition measure (100-120 ms) |  |  |  |  |
| *Intercept* | 0.9657 | 0.1109 | 8.712 | <0.0001 |
| SeqFwd ‘Other’ Tx | -0.1115 | 0.0812 | -1.373 | 0.1480 |
| SeqBkw ‘Other’ Tx | -0.0013 | 0.0802 | -0.016 | 0.9712 |
| **After condition:** current + other transition (100-120 ms) |  |  |  |  |
| *Intercept* | 0.8549 | 0.1127 | 7.589 | <0.0001 |
| SeqFwd | -0.1268 | 0.0853 | -1.488 | 0.1248 |
| SeqBkw | 0.1540 | 0.0823 | 1.872 | 0.0472* |
| SeqFwd ‘Other’ Tx | 0.1053 | 0.0850 | 1.239 | 0.2320 |
| SeqBkw ‘Other’ Tx | -0.0176 | 0.0870 | -0.203 | 0.8368 |
| **Before condition:** current + other transition (100-120 ms) |  |  |  |  |
| *Intercept* | 0.9728 | 0.1128 | 8.627 | <0.0001 |
| SeqFwd | 0.1796 | 0.0782 | 2.296 | 0.0160* |
| SeqBkw | -0.0153 | 0.0790 | -0.194 | 0.8352 |
| SeqFwd ‘Other’ Tx | -0.1346 | 0.0862 | -1.562 | 0.1240 |
| SeqBkw ‘Other’ Tx | -0.0745 | 0.0865 | -0.861 | 0.3536 |

**Table S6.** Multilevel modeling results relating cue-evoked responses (200 – 250 ms post-onset) and sequenceness to accuracy. Top: inclusion of cue-evoked representation of current episode categories (omitting the on-screen category) minus other-episode categories. Bottom: inclusion of the response to the cue category itself.

| **After condition: Current-other episode reactivation** |  |  |  |  |
| --- | --- | --- | --- | --- |
| **Accuracy ~** | **coef.** | **ste** | **z-stat** | **p-value** |
| *Intercept* | 0.8534 | 0.1128 | 7.562 | <0.0001 |
| SeqFwd | -0.1366 | 0.0779 | -1.753 | 0.0816 |
| SeqBkw | 0.1862 | 0.0779 | 2.39 | 0.0160* |
| Current-Other coef. | 0.8489 | 0.6786 | 1.251 | 0.2040 |
| **Before condition: Current-other episode reactivation** |  |  |  |  |
| *Intercept* | 0.962 | 0.112 | 8.586 | <0.0001 |
| SeqFwd | 0.1601 | 0.0727 | 2.202 | 0.0256* |
| SeqBkw | -0.0574 | 0.0739 | -0.777 | 0.4240 |
| Current-Other coef. | 0.5279 | 0.7212 | 0.732 | 0.4400 |
| **After condition: Cued category response** |  |  |  |  |
| *Intercept* | 0.847 | 0.1123 | 7.541 | <0.0001 |
| SeqFwd | -0.1276 | 0.078 | -1.636 | 0.0984 |
| SeqBkw | 0.1882 | 0.0779 | 2.415 | 0.018* |
| Cue response coef. | 1.5659 | 0.5896 | 2.656 | 0.0016* |
| **Before condition: Cued category response** |  |  |  |  |
| *Intercept* | 0.9615 | 0.1118 | 8.597 | <0.0001 |
| SeqFwd | 0.16 | 0.0727 | 2.202 | 0.0256* |
| SeqBkw | -0.0564 | 0.074 | -0.763 | 0.4464 |
| Cue response coef. | 0.0133 | 0.7267 | 0.018 | 0.8424 |

**Table S7.** Multilevel modeling results for the interaction between sequenceness and episode length (long, short) or episode end valence (positive, negative) on accuracy.

| **After condition: Length** |  |  |  |  |
| --- | --- | --- | --- | --- |
| **Accuracy ~** | **coef.** | **ste** | **z-stat** | **p-value** |
| *Intercept* | 0.8415 | 0.1129 | 7.456 | <0.0001 |
| SeqFwd | -0.1375 | 0.0782 | -1.757 | 0.0808 |
| SeqBkw | 0.1948 | 0.0783 | 2.487 | 0.0080* |
| Length | -0.0909 | 0.0822 | -1.105 | 0.2912 |
| SeqFwd * Length | 0.013 | 0.0783 | 0.166 | 0.8680 |
| SeqBkw * Length | 0.0105 | 0.0786 | 0.134 | 0.8938 |
| **Before condition: Length** |  |  |  |  |
| *Intercept* | 0.9602 | 0.1127 | 8.517 | <0.0001 |
| SeqFwd | 0.1612 | 0.0732 | 2.204 | 0.0296* |
| SeqBkw | -0.0616 | 0.0741 | -0.831 | 0.4152 |
| Length | 0.0744 | 0.08 | 0.929 | 0.3672 |
| SeqFwd * Length | -0.0849 | 0.0732 | -1.159 | 0.2664 |
| SeqBkw * Length | 0.0529 | 0.0744 | 0.711 | 0.4728 |
| **After condition: End valence** |  |  |  |  |
| *Intercept* | 0.8702 | 0.1166 | 7.463 | <0.0001 |
| SeqFwd | -0.1175 | 0.079 | -1.488 | 0.1208 |
| SeqBkw | 0.1821 | 0.0789 | 2.307 | 0.0200* |
| Reward | -0.0873 | 0.0842 | -1.037 | 0.3000 |
| SeqFwd * Valence | 0.1108 | 0.079 | 1.402 | 0.1760 |
| SeqBkw * Valence | -0.2005 | 0.08 | -2.505 | 0.0128* |
| **Before condition: End valence** |  |  |  |  |
| *Intercept* | 0.964 | 0.1116 | 8.638 | <0.0001 |
| SeqFwd | 0.1561 | 0.0733 | 2.129 | 0.0304* |
| SeqBkw | -0.0541 | 0.0744 | -0.726 | 0.4624 |
| Reward | -0.005 | 0.0915 | -0.055 | 0.9648 |
| SeqFwd * Valence | -0.0071 | 0.0734 | -0.097 | 0.9104 |
| SeqBkw * Valence | -0.0047 | 0.0745 | -0.062 | 0.9344 |

**Table S8.**
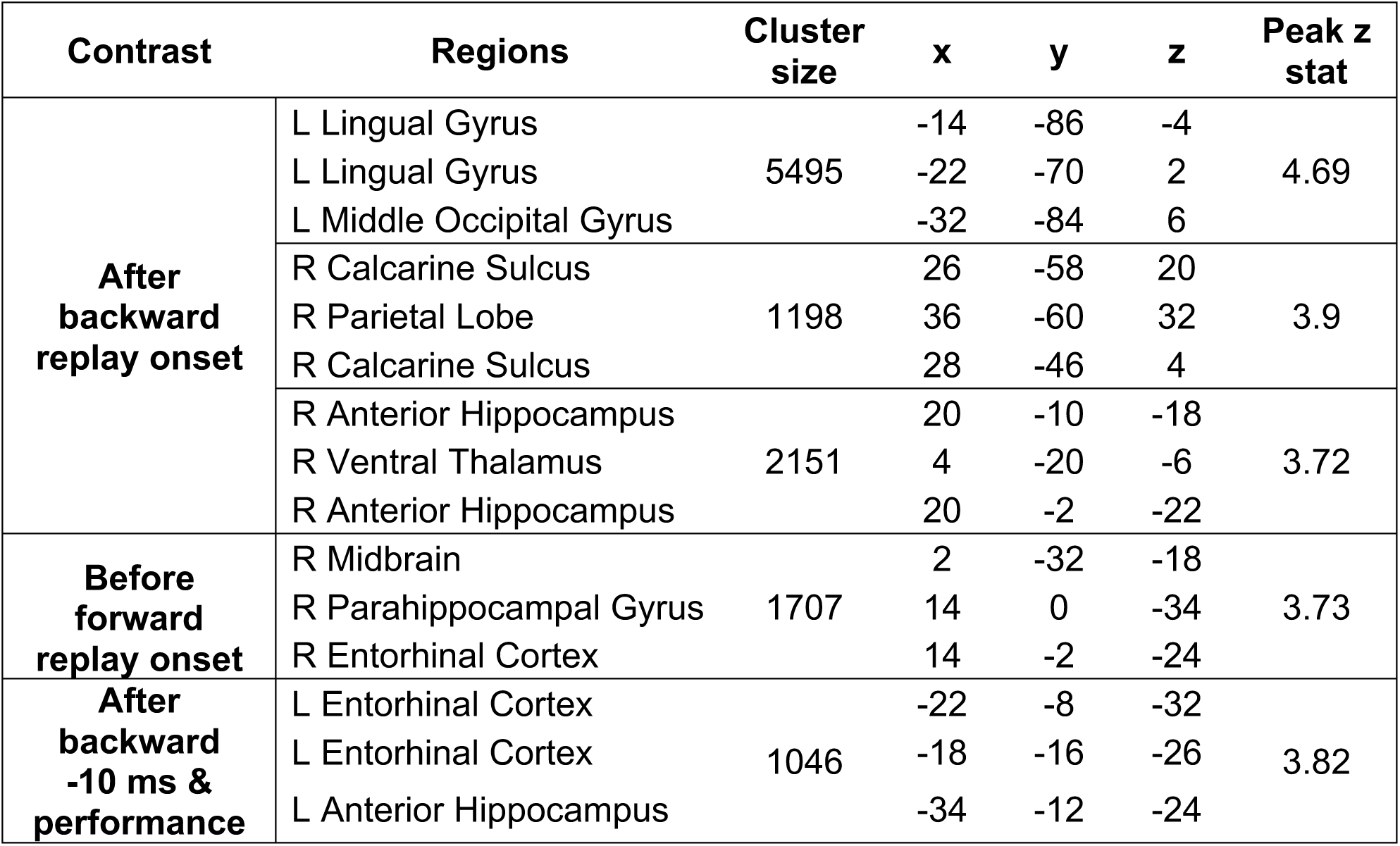
Whole-brain beamforming MEG results for replay onset in the after and before conditions. Clusters significant whole-brain FWE-corrected after an initial threshold of p < 0.001 to provide interpretable clusters.

